# Proteolytic feedback of quorum sensing promotes infection by *Pseudomonas aeruginosa*

**DOI:** 10.64898/2026.11.02.763748

**Authors:** Caleb P. Mallery, Sheryl E. Fernandes, Megan L. Schumacher, Autumn N. Pope, Hayden Dillow, Alicia G. Mendoza, Peter A. Jorth, Jon E. Paczkowski

## Abstract

Quorum sensing (QS) enables bacteria to coordinate collective behaviors in response to population density. However, the mechanisms by which QS circuits are reset or attenuated remain poorly understood. In *Pseudomonas aeruginosa*, the LuxR-type transcription factor RhlR controls virulence gene expression at high cell density, requiring both its cognate acyl-homoserine lactone signal and the accessory protein PqsE for full activity. Here, we show that RhlR abundance is actively controlled by ATP-dependent proteolysis through a self-limiting negative feedback loop. We identify an N-terminal degron within RhlR that is protected from protease recognition by PqsE binding and is required for degradation by both the Lon and the ClpP1-ClpP2 protease complex. Removal of this degron stabilizes RhlR, advancing the timing of QS-regulated gene expression, enhancing biofilm formation, imposes a competitive fitness cost *in vitro*, and reduced virulence in an acute mouse lung infection. Mechanistically, RhlR directly activates transcription of *clpP2*, encoding a subunit of the protease complex required for its own degradation, establishing a self-limiting negative feedback loop. Disruption of RhlR DNA binding stabilizes the receptor through dual effects on reduced *clpP2* expression and increased PqsE availability via the loss of RhlR- dependent repression of *pqsE*. Together, these findings reveal proteolytic turnover as an underappreciated regulatory layer of QS-activation and establish a mechanism for how receptor turnover may facilitate the transition away from high-cell density states.

## INTRODUCTION

*Pseudomonas aeruginosa* is an opportunistic pathogen of growing global concern, causing life-threatening infections in immunocompromised individuals and posing an increasing threat due to its multi-drug resistance and persistence in hospital settings^1,2^. Central to its pathogenicity is quorum sensing (QS), a mechanism of bacterial communication that allows bacteria to sense and respond to the population density of their environment^3–5^. Through the production and detection of small diffusible chemical signals, bacteria coordinate gene expression across the population enabling collective behaviors that would be unachievable by a single cell. Group behaviors include biofilm formation, toxin production, and secretion, which collectively shape how bacteria interact with their environments and hosts^6^. Understanding the molecular mechanisms that regulate QS is therefore central to understanding bacterial pathogenesis.

In Gram-negative bacteria, QS is often mediated by acyl-homoserine lactone (AHL) signals. AHLs are synthesized by LuxI-family enzymes and accumulate in the extracellular environment as population density increases, freely diffusing across membranes to reach intracellular cytoplasmic receptors^7–9^. These receptors, known as LuxR-type transcription factors, bind AHLs and modulate expression of QS-regulated target genes as well as the *luxI* synthase gene, creating a positive feedback loop that amplifies signaling as the population increases^3,9,10^.

*P. aeruginosa* employs multiple interconnected QS circuits including the AHL based Las and Rhl systems and the non AHL, *Pseudomonas* Quinolone Signal (PQS) system^11^. Among these circuits, the Rhl system plays a central role in controlling virulence factor expression. The LuxR-type transcription factor RhlR activates genes required for pyocyanin production, rhamnolipid biosynthesis, and biofilm maturation, among others^12–17^. Unlike many LuxR-type receptors, RhlR requires high concentrations of its cognate AHL signal *N*-butanoyl homoserine lactone (C4-HSL) for half-maximal activation (EC50) suggesting that robust Rhl signaling is tightly regulated^18,19^. Notably, RhlR function is also modulated by PqsE, a thioesterase in the PQS system. Although initially characterized for its enzymatic role in 2-alkyl-4-quinolone biosynthesis^20^, PqsE plays an additional independent role in the Rhl system through direct physical interaction with RhlR^21^. This interaction is separable from PqsE enzymatic activity, and modulates RhlR-dependent gene expression^16,22–26^.

The ability of bacteria to rapidly coordinate their gene expression not only depends on the production of transcriptional regulators, but also their timely removal. Selective protein degradation enables the precise timing of gene expression and the prevention of inappropriate activation^27^. In bacteria, this regulatory proteolysis is carried out by ATP-dependent AAA+ (**A**TPases **a**ssociated with cellular **a**ctivities) proteases, including Lon and the Clp family, among others28. These proteases display distinct substrate preferences and are responsible for substrate unfolding and threading them into a proteolytic chamber for degradation. AAA+ proteases can be broadly divided into two functional components: an AAA+ unfoldase, which recognizes protein substrates and performs the initial unfolding, and a peptidase, which performs the proteolytic cleavage (**Fig. 1a**). In some cases, such as Lon, both components are encoded within a single multifunctional protein^29^. In others, such as the Clp family, unfolding and peptidase activities are carried out by separate proteins that associate to form a hetero-oligomeric complex^30^. For example, ClpP is a tetradecameric (14-subunit) peptidase that can associate with hexamers (6-subunit) of the unfoldase ClpX6 to form ClpX6P14, or with ClpA6 to form ClpA6P14, each with distinct substrate specificities (**Fig. 1a**)^31^. AAA+ proteases are appreciated as broad regulators, targeting diverse regulatory proteins to tune gene expression in response to environmental and physiological cues^27,32,33^. Given that LuxR-type receptors govern QS across a wide range of Gram-negative pathogens, understanding how their abundance is post-translationally controlled has broad relevance^10,34,35^. Accumulating evidence indicates that proteolysis shapes QS receptor abundance and activity^36,37^, yet the precise mechanisms remain poorly defined for *P. aeruginosa* and LuxR-type receptors, in general.

**Fig. 1:**
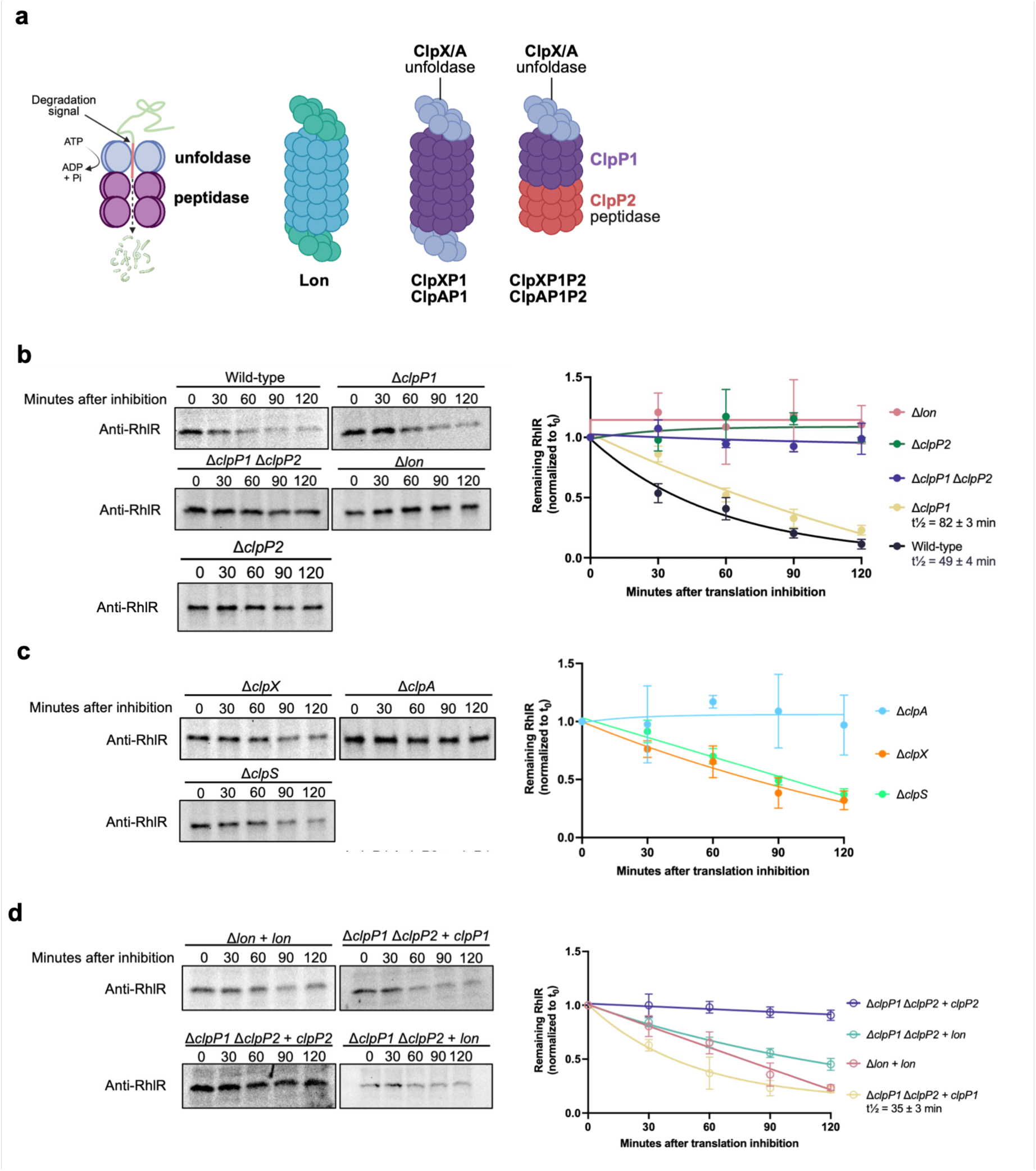
Lon and Clp-proteases regulate RhlR turnover and steady-state abundance. a, (Left) Simplified mechanism of AAA+ proteases. A hexameric ring (blue) recognizes a degradation signal on a protein substrate, then uses ATP binding and hydrolysis to unfold and translocate it into a peptidase chamber (purple) for degradation. (Right) domain organization of the proteases discussed here. In lon, the unfoldase and peptidase domains are fused within a single polypeptide, forming a homo-oligomeric complex. Clp proteases instead assemble from a barrel-shaped peptidase core, built from two stacked heptameric rings, that is activated by docking of a hexameric ClpX or ClpA unfoldase on one or both faces. ClpP1 can form a homomeric core (ClpP114) singly or doubly capped by ClpX/ClpA, whereas ClpP1-ClpP2 assemble into a heteromeric core (ClpP17P27) that can only be singly capped, with ClpX/ClpA docking on the ClpP1 face. **b**, RhlR degradation following translation inhibition in protease mutants. Wild-type (WT), Δ*clpP1*, Δ*clpP1*/Δ*clpP2*, and Δ*lon* strains were grown to OD600 ∼2.0, and translation was inhibited. Samples were collected at the indicated times and analyzed by immunoblotting with anti-RhlR antibody. Quantification of RhlR remaining over time (normalized to t0) is shown on the right. Data represent mean ± s.d. of three independent biological replicates. Apparent half-lives were calculated by exponential decay fitting. **c,** RhlR degradation following translation inhibition in protease unfoldase/adaptor protein mutants. **d**, Complementation analysis of protease requirements for RhlR degradation. Δ*lon* + lon, Δ*clpP1* Δ*clpP2* + *clpP1*, Δ*clpP1* Δ*clpP2* + *clpP2*, and Δ*clpP1* Δ*clpP2* + *lon* strains were subjected to translation inhibition as in (a). RhlR levels were quantified relative to t0. Data represent mean ± s.d. of three biological replicates.

A well-characterized example of protease-dependent QS regulation involves LuxR-type receptors themselves, many of which are intrinsically unstable in the absence of their cognate AHL ligand. The paradigmatic receptor TraR fails to fold properly without AHL; thus, ligand binding protects the protein from degradation by Lon and Clp proteases^38^. In *P. aeruginosa*, both Lon and Clp proteases have been implicated in QS regulation. *lon* mutants display impaired motility, defective biofilm formation, and reduced pyocyanin production^39–41^, while ClpXP-mediated proteolysis suppresses QS by degrading key components of the PQS and Rhl signaling pathways^42–46^. Notably, *P. aeruginosa* encodes two ClpP paralogs with distinct features^47^. ClpP1 is constitutively expressed and assembles into an active 14-subunit homopeptidase. In contrast, ClpP2 is expressed during stationary phase under QS control^46^ with both LasR and RhlR binding sites identified upstream in the *clpP2* promoter^17,48^. Importantly, ClpP2 is not functional on its own as it forms inactive heptamers (7-subunit) ClpP2749. However, ClpP2 is active upon hetero-oligomerization with ClpP1, forming an active protease complex in which one heptameric ring of ClpP1 stacks onto a heptameric ring of ClpP2, forming a ClpP17ClpP27 complex (**Fig. 1a**)^47,49,50^.

Contrary to the canonical model, previous work from our group demonstrated that RhlR stability is not dependent on its cognate AHL signal. Deletion of *rhlI*, the synthase responsible for C4-HSL production, did not impair RhlR abundance *in vivo*^37,51^. Instead, RhlR stability is governed by the co-regulator PqsE. Deletion of *pqsE* or disruption of its ability to interact with RhlR decreased RhlR abundance. Deletion of *lon* increased RhlR levels. However, a Δ*pqsE* Δ*lon* strain did not exhibit restored RhlR levels, implicating additional proteases beyond Lon in RhlR turnover^37^. Together, these findings suggest that RhlR stability is controlled through an unconventional signal-independent mechanism and prompted us to investigate the proteolytic mechanism responsible for RhlR turnover.

Here, we show that RhlR is a target of ATP-dependent proteolysis by both Lon and Clp proteases. We identify an N-terminal degron (residues 1–17) required for RhlR degradation and demonstrate that removal of this degron (ΔN17 RhlR) stabilizes RhlR *in vivo*, even in the absence of *pqsE* protection. Stabilized RhlR results in elevated protein levels and accelerates the timing of QS gene expression, including enhanced biofilm formation and increased production of QS-dependent metabolites during early growth. Importantly, stabilization of RhlR imposes a measurable fitness cost under multiple growth conditions, and this cost is fully suppressed in a Δ*rhlR* background, indicating that regulated RhlR turnover is essential for competitive fitness. Mechanistically, we establish that PqsE protects the N-terminal degron of RhlR from degradation while RhlR directly activates *clpP2* transcription, encoding a peptidase required for its own degradation, establishing a negative feedback loop that limits RhlR abundance. Together, these findings uncover an unappreciated protease-mediated layer of QS regulation in *P. aeruginosa* and highlight a self-limiting mechanism of RhlR control that fine tunes QS outputs.

## RESULTS

### Multiple ATP-dependent proteases contribute to RhlR turnover

Recent work from our group demonstrated that the QS regulator RhlR is degraded in the absence of *pqsE* by Lon protease^37^. To further define the proteolytic systems responsible for RhlR turnover, we monitored RhlR stability following translation inhibition in strains lacking individual ATP-dependent proteases. Protease deletions did not impair growth (**Supplementary Fig. 1a, b**), and pyocyanin production was largely unaffected except in Lon-deficient backgrounds (**Supplementary Fig. 1c,d**), which showed reduced pyocyanin, consistent with prior reports^40^. To determine the individual role of different components of the proteasome, we monitored RhlR protein levels after stopping translation with the addition of chloramphenicol. In wild-type cells, RhlR levels decreased steadily over time, exhibiting a half-life of approximately 49 minutes (**Fig. 1b**). Deletion of *clpP1* partially stabilized RhlR and increased its half-life to approximately 82 minutes while deletion of both *clpP1* and *clpP2* markedly stabilized RhlR (**Fig. 1b**). Notably, deletion of *clpP2* alone fully stabilized RhlR despite ClpP2 having no independent proteolytic activity (**Fig. 1b**). This paradox suggests that ClpP2 contributes substrate specificity to the heterocomplex. The presence of ClpP2 is likely required to direct ClpP1-ClpP2 activity toward RhlR, even though it cannot degrade substrates independently^47,49^. Consistent with our previous findings^37^, deletion of *lon* also stabilized RhlR to levels comparable to that of a double deletion of *clpP1/P2* (**Fig. 1b**). Together, these data demonstrate that both the ClpP1-ClpP2 complex and Lon contribute to RhlR turnover.

To identify the ATPase components that contribute to ClpP-dependent degradation of RhlR, we next examined mutants lacking individual unfoldases or adaptor proteins. Deletion of *clpA* reproducibly stabilized RhlR relative to wild-type cells, whereas deletion of *clpX* produced only modest effects under the tested conditions (**Fig. 1c**). In contrast, deletion of *clpS* did not appreciably alter RhlR turnover (**Fig. 1c**), suggesting that degradation does not proceed through a canonical ClpS-dependent N-end rule pathway^52^. These findings indicate that ClpA contributes substantially to RhlR degradation, although the partial effects observed across multiple mutants suggest that RhlR may be recognized by more than one ATP-dependent proteolytic pathway.

To confirm the contributions of these proteolytic systems, we complemented the respective deletion strains. As expected, expression of *lon* restored RhlR turnover in the Δ*lon* mutant background while expression of *clpP1* restored degradation in the Δ*clpP1* mutant to near wild-type levels (t½ ≈ 35 min; **Fig**. **1d**). In contrast, expression of *clpP2* alone in the Δ*clpP1* Δ*clpP2* background did not restore detectable RhlR turnover, consistent with the inability of ClpP2 to function as an active protease independently of ClpP1 (**Fig. 1d**). Interestingly, overexpression of *lon* in the Δ*clpP1* Δ*clpP2* mutant promoted RhlR degradation despite the absence of ClpP1-ClpP2 activity (**Fig. 1d**), indicating that elevated Lon activity can partially compensate for loss of Clp-mediated proteolysis. The inability of endogenous Lon to fully compensate for the loss of ClpP1-ClpP2 activity, despite partial restoration of RhlR turnover upon *lon* overexpression, suggests that these proteolytic systems contribute non-equivalently to RhlR degradation. Together, these findings suggest that multiple ATP-dependent proteolytic systems can recognize RhlR, potentially recognizing distinct conformational or regulatory states of the protein under different cellular conditions.

### RhlR directly activates expression of the *clpP2* peptidase

Having established that Clp protease contributes to RhlR turnover, we asked whether QS circuitry itself regulates expression of *clpP2*. Multiple lines of evidence have suggested that the protease gene *clpP2* is controlled by QS^17,53–55^. Previous ChIP-seq studies identified a RhlR binding site upstream of *clpP2*^17^, while EMSA experiments demonstrated that LasR can bind the same promoter region *in vitro*^48^. In addition, RNA-seq analyses reported reduced *clpP2* expression in *lasR* mutant backgrounds, raising the possibility that LasR directly activates *clpP2* transcription^56^. Because LasR functions upstream of RhlR in the canonical QS hierarchy, it remained unclear whether LasR directly activates *clpP2* or instead regulates *clpP2* indirectly through RhlR.

To distinguish between these possibilities, we tested *clpP2* activation using a heterologous *Escherichia coli* reporter system carrying a p*clpP2*-*luxCDABE* fusion. Expression of RhlR strongly activated p*clpP2* in the presence of C4-HSL (**Fig. 2a**). Prior work showed that PqsE attenuates RhlR activation of the *clpP2* promoter^22^, prompting us to test whether constitutive expression of *pqsE* attenuated RhlR-dependent expression here, which it did (**Fig. 2a**). However, in *P. aeruginosa*, PqsE did not impact *clpP2* levels as shown by RT-qPCR in *pqsE* deletion and overexpression strains grown to stationary phase, compared to wild type (**Supplementary Fig. 2**), indicating that the *E. coli* attenuation does not reflect the native context.

**Fig. 2:**
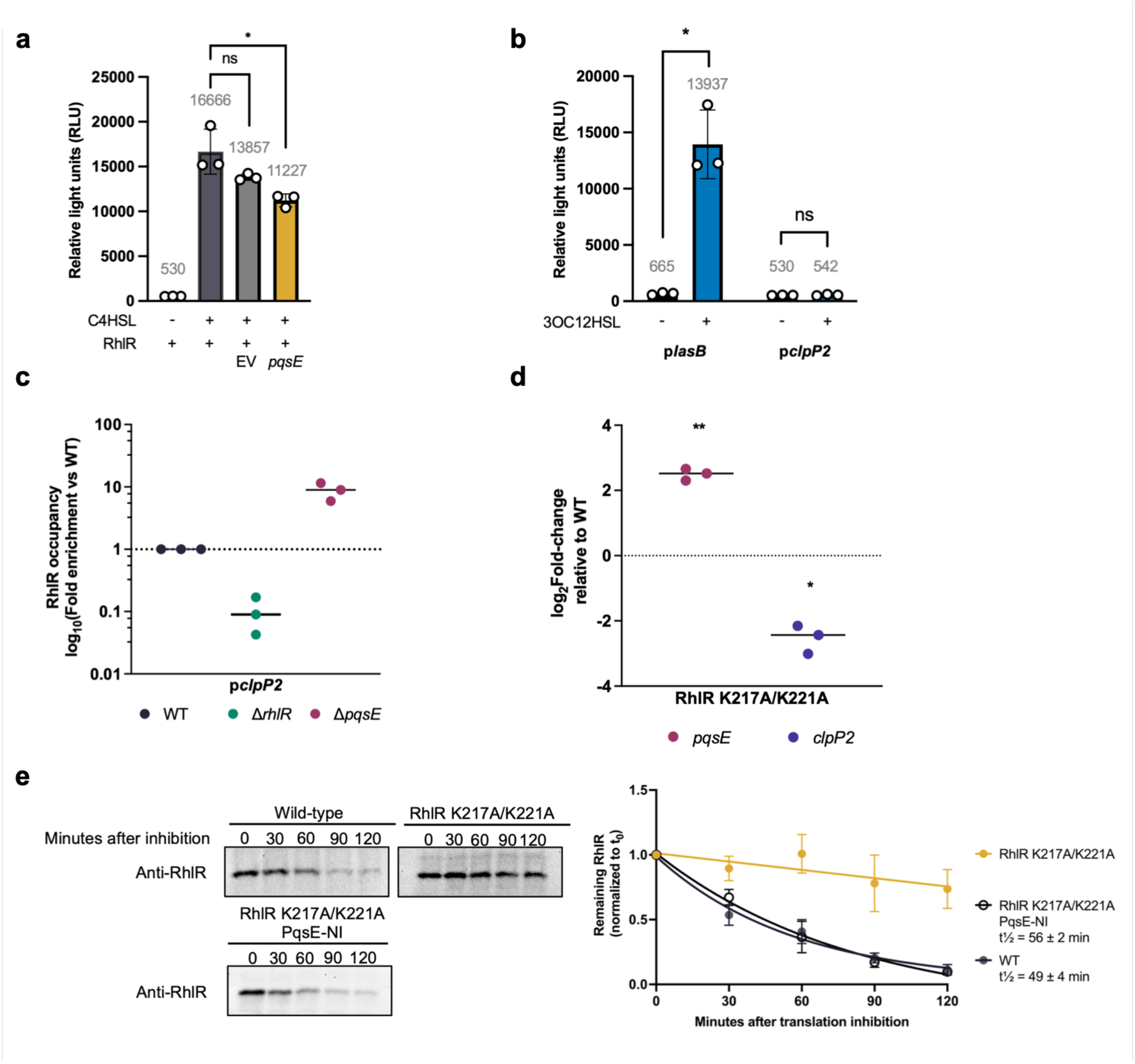
RhlR directly activates *clpP2* expression through DNA binding and modulates its own stability. **a**, Activation of p*clpP2-lux* in *E. coli* is reported as relative light units with RhlR expressed from an arabinose-inducible promoter in the presence or absence of 10 µM C4-HSL and co-expressed with empty vector (EV) or *pqsE*. Data represent mean ± SD of three independent biological replicates. Significance was determined by unpaired t-test; *p<0.05; ns, not significant. **b**, Activation of p*clpP2-luxCDABE* in *E. coli* is reported as relative light units with LasR expressed from an arabinose inducible promoter in the presence or absence of 10 µM 3OC12-HSL. Data represent mean ± SD of three independent biological replicates. Significance was determined by unpaired t-test; \**P* < 0.05; ns, not significant. **c**, Direct chromatin immunoprecipitation analysis of RhlR occupancy at the *clpP2* promoter in various genetic backgrounds. RhlR occupancy is represented as log10 fold-enrichment relative to WT which is normalized to 1. **d**, *pqsE* and *clpP2* transcript levels were measured by RT-qPCR in the strain expressing the RhlR K217A/K221A mutant relative to WT. Data are shown as log2 fold-change relative to WT (2^-ΔΔCt^). Each point represents one biological replicate (n = 3), with the horizontal line indicating the mean. Statistical significance was determined by unpaired two-tailed Student’s t-test with Welch’s correction on ΔCt values for each gene relative to WT. \**P* < 0.05, \*\**P* < 0.01. **e**, RhlR degradation following translation inhibition in DNA-binding mutants. Top left panel shows a representative Western blot of RhlR protein levels over time. Bottom panel displays quantified RhlR degradation curves for WT (t½ ≈ 49 ± 4 minutes), RhlR K217A/K221A, and RhlR K217A/K221A + PqsE-NI (t½ ≈ 56 ± 2 minutes). Apparent half-lives were calculated by exponential decay fitting.

To determine whether LasR directly activates *clpP2*, we tested LasR in the heterologous reporter system. LasR did not activate *p*clpP2 (**Fig. 2b**). For comparison, LasR with N-(3-oxododecanoyl)-L-homoserine lactone (3OC12-HSL) strongly activated p*lasB*, confirming that the reporter system was responsive to LasR. However, RT-qPCR in *P. aeruginosa* revealed an indirect role for LasR; a Δ*lasR* mutant showed decreased *clpP2* expression (**Supplementary Fig. 3**), consistent with previous studies^56^. As expected for an established activator, a Δ*rhlR* mutant also showed decreased *clpP2* expression (**Supplementary Fig. 3**). Notably, overexpression of *rhlR* in the Δ*lasR* background restored *clpP2* expression to wild-type levels, demonstrating that LasR controls *clpP2* indirectly through its positive regulation of *rhlR* expression, rather than direct promoter activation (**Supplementary Fig. 3**).

Direct chromatin immunoprecipitation (ChIP) qPCR performed using *P. aeruginosa* cells confirmed that RhlR directly binds the *clpP2* promoter *in vivo* (**Fig. 2c**). Binding was ablated in a Δ*rhlR* mutant, confirming RhlR specificity. Importantly, RhlR binding was enriched approximately 7-10-fold in the Δ*pqsE* mutant relative to wild-type, demonstrating that the presence of PqsE attenuates RhlR occupancy at the *clpP2* promoter (**Fig. 2c**). This attenuation of RhlR DNA binding is consistent with the reduced p*clpP2* activation observed upon constitutive *pqsE* expression in the *E. coli* reporter (**Fig. 2a**). However, despite the increased RhlR occupancy in the absence of *pqsE*, *clpP2* transcript levels were unchanged in *P. aeruginosa* strains lacking or overexpressing *pqsE* (**Supplementary Fig. 2**). Thus, PqsE modulates RhlR occupancy at the *clpP2* promoter without substantially altering *clpP2* transcription in the native conditions tested.

Thus far, we have established that RhlR directly activates *clpP2*, a protease component involved in RhlR degradation, while simultaneously repressing *pqsE*, which encodes a RhlR stabilizer. This creates a potential self-limiting circuit; RhlR activates the protease machinery for its own degradation while suppressing PqsE, which protects it from degradation. To test whether RhlR actually initiates this self-limiting feedback, we examined the stability of a RhlR DNA binding mutant, RhlR K217A/K221A which cannot activate *clpP2* or repress *pqsE*^57–59^. If the feedback loop is functional, we hypothesize that this mutant will show decreased *clpP2* and elevated *pqsE* expression leading to RhlR protein stability in our DNA binding mutant. RT-qPCR confirmed this hypothesis (**Fig. 2d**), RhlR K217A/K221A showed reduced *clpP2* and elevated *pqsE* transcripts relative to wild-type, establishing that RhlR directly controls *clpP2* expression through DNA binding and simultaneously de-represses *pqsE* expression when DNA binding is abolished. RhlR protein stability correlated with these data (**Fig. 2d**). As shown previously, wild-type RhlR was degraded with a half-life of approximately 49 minutes, whereas the RhlR K217A/K221A variant was stable across the time course (**Fig. 2e**), reflecting reduced *clpP2* and increased *pqsE* levels. To test whether the PqsE-RhlR interaction is required for the observed stability, we examined the RhlR K217A/K221A variant in combination with a non-interacting PqsE variant (PqsE-NI) defective in RhlR binding^16^. This strain showed decreased RhlR levels back to wild-type levels, confirming that the PqsE-RhlR interaction is necessary for RhlR protection from degradation. Together, these results confirm that *clpP2* is a target of RhlR and that RhlR regulates its own stability through a self-limiting mechanism dependent on both ClpP2-mediated degradation and PqsE protection.

### An N-terminal degron mediates RhlR stability *in P. aeruginosa*

Because deletion of *pqsE* destabilizes RhlR, we reasoned that PqsE may protect a protease recognition element within RhlR. Proteolytic degradation of LuxR-type transcription factors or similar family of receptors has been described in a limited number of systems^38,60^, and the sequence determinants governing receptor turnover remain poorly understood. Inspection of our recently solved PqsE-RhlR structure identified the N-terminus of RhlR (residues 1-17) as a solvent exposed region positioned at the binding interface of PqsE dimers (**Fig. 3a**; dark green inset). Notably, this region contains a cluster of hydrophobic and aromatic residues (FLLWWD), which could serve as a recognition element for AAA+ proteases^61,62^. We therefore hypothesized that residues 1–17 function as a degron with accessibility modulated by PqsE binding and tested this by replacing the native *rhlR* allele with a truncation lacking the N-terminal 17 amino acids (ΔN17 RhlR). To assess whether this region is required for turnover, we compared RhlR stability across wild-type, ΔN17 RhlR, and Δ*pqsE* backgrounds following translation inhibition. As before, following translation inhibition, wild-type RhlR exhibited a half-life of approximately 49 minutes (**Fig. 3b**). Consistent with our previous observations, deletion of *pqsE* accelerated RhlR turnover by approximately 10 minutes. The Δ*pqsE* strain started with lower steady-state RhlR levels due to the loss of PqsE-mediated stabilization^37^; however, despite this reduced pool, RhlR was degraded at a faster rate than the wild-type background, further supporting a protective role for PqsE (**Fig. 3b**). In contrast, ΔN17 RhlR remained stable throughout the 120-minute time course. Importantly, deletion of *pqsE* failed to destabilize ΔN17 RhlR, demonstrating that the N-terminus of RhlR is required for turnover in the absence of *pqsE*. Together, these findings identify the N-terminus of RhlR as a critical determinant of proteolytic turnover and establish a link between PqsE-dependent stabilization and protease recognition.

**Fig. 3:**
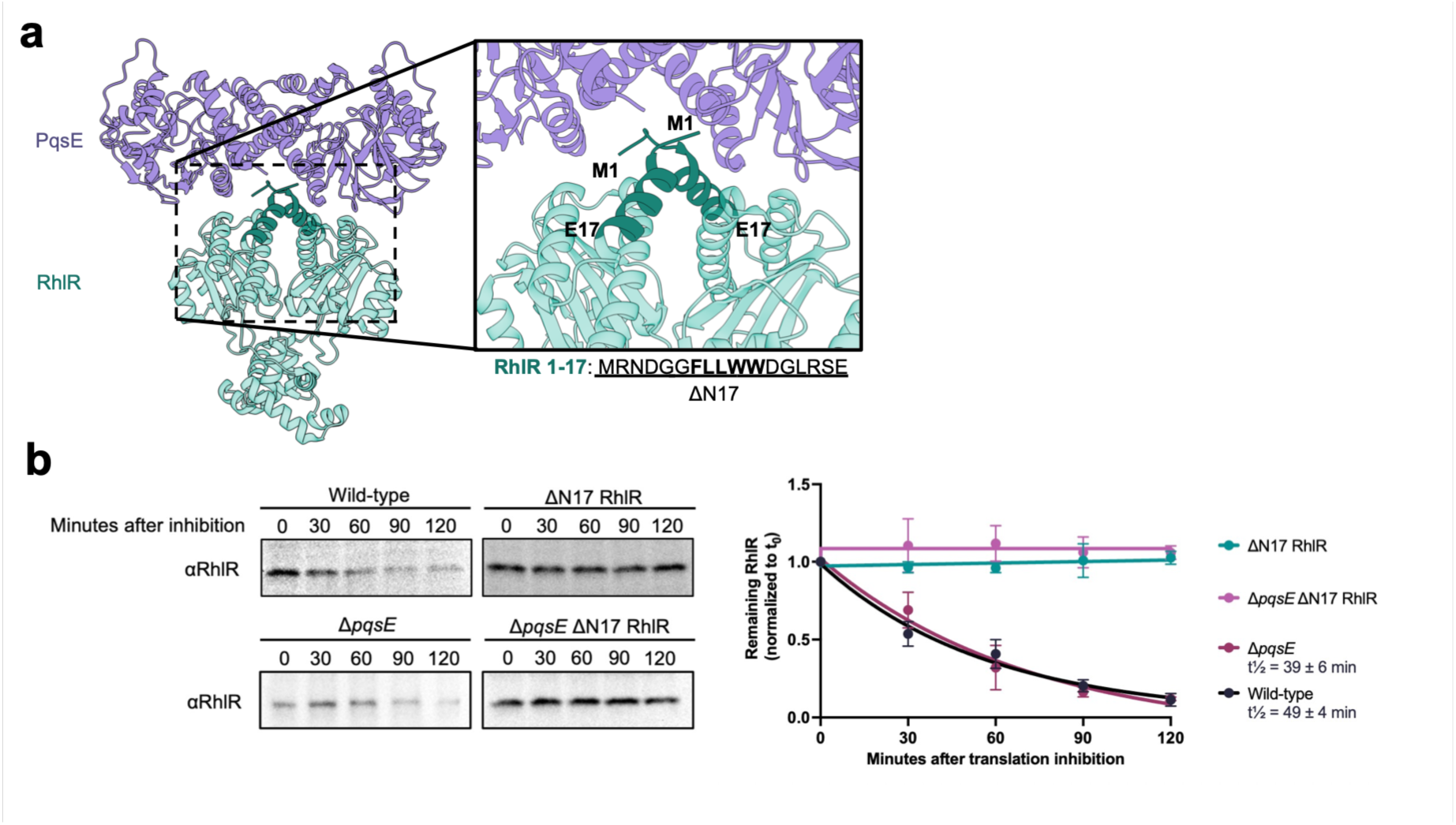
An N-terminal degron confers RhlR stability *in P. aeruginosa*. **a**, Structural model of the PqsE-RhlR complex (PDB ID: 8DQ0^59^). RhlR (teal) and PqsE (purple) are shown as ribbon representations. Inset highlights the RhlR N-terminus (residues 1-17), predicted to be flexible and solvent exposed in the absence of PqsE are colored in dark green. The N-terminal sequence (residues 1-17) is shown below, with hydrophobic residues in bold. **b**, RhlR degradation following translation inhibition. Wild-type (WT), Δ*pqsE*, ΔN17 RhlR, and *pqsE*/ΔN17 RhlR strains were grown to an OD600 of ∼2.0, and translation was inhibited. Samples were collected at the indicated times and analyzed by immunoblotting with anti-RhlR antibodies. Quantification of remaining RhlR over time (normalized to t0) is shown on the right. Data represent the mean ± s.d. of three biological replicates. Apparent half-lives were calculated by exponential decay fitting and displayed for strains where one could be detected.

Because residues 1-17 are positioned near the PqsE-RhlR interface, we next asked whether the truncation disrupted the functional interaction between these proteins. Pyocyanin production requires formation of the PqsE-RhlR complex^16^, providing a convenient readout of the interaction. ΔN17 RhlR produced pyocyanin at levels comparable to wild type after overnight planktonic growth, suggesting that deletion of the N-terminal region does not abolish PqsE binding (**Supplementary Fig. 4a**). Thus, stabilization of ΔN17 RhlR is unlikely to arise from disruption of the PqsE-RhlR interaction. We attempted to biochemically characterize ΔN17 RhlR; however, heterologous expression of ΔN17 RhlR in *E. coli* yielded insoluble protein and failed to activate a transcriptional reporter (**Supplementary Fig. 4b**), preventing analysis outside the native *P. aeruginosa* background. Together, these data identify residues 1-17 as an N-terminal degron required for RhlR turnover *in vivo*. We propose that interaction with PqsE limits access of this degron, thereby protecting RhlR from degradation by Lon and Clp-dependent proteolysis (**Fig. 3d**).

### RhlR steady-state abundance is controlled by proteolysis and PqsE

The above experiments identified an N-terminal degron within RhlR and established that Lon and ClpP1-ClpP2 promote RhlR degradation while PqsE protects RhlR from turnover. We next asked how these regulatory pathways influence RhlR abundance under physiological growth. To assess RhlR steady-state levels, cells were harvested after 6 hours of growth, corresponding to the zero timepoint of the translation inhibition assays. Relative to wild-type, ΔN17 RhlR, Δ*lon*, and Δ*clpP1* Δ*clpP2* strains accumulated 5- to 8-fold more RhlR (**Fig. 4a**), demonstrating that both protease systems and the N-terminal degron constrain RhlR abundance during normal growth.

**Fig. 4:**
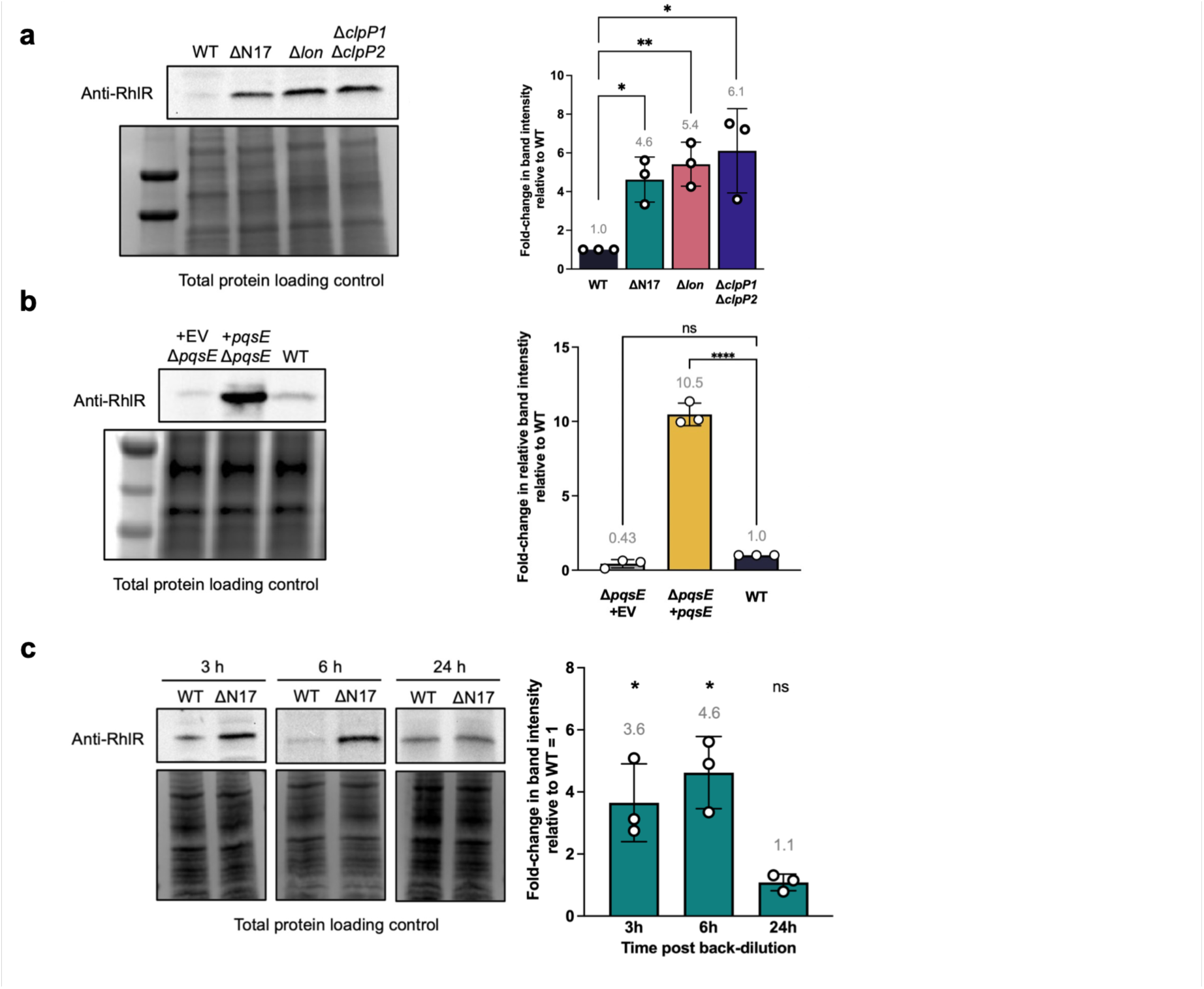
RhlR steady-state abundance is controlled by proteolysis and PqsE. **a**, Steady-state RhlR abundance in WT PA14 and protease-deficient strains. Cells were harvested at OD600 ∼2.0 and analyzed by immunoblotting. Band intensities were normalized to total protein staining and shown relative to WT. Representative blot (left) and quantification (right) are shown. Data represent mean ± s.d. of three independent biological replicates. **b**, Effect of PqsE and protease deletions on RhlR steady-state levels. RhlR abundance was measured in Δ*pqsE* strains carrying empty vector (EV) or constitutively expressed *pqsE*, as well as in Δ*pqsE* double- and triple-mutants (Δ*pqsE* ΔN17 RhlR, ΔpqsE Δ*lon*, and Δ*pqsE* Δ*clpP1* Δ*clpP2*). Band intensities were normalized to total protein staining and expressed relative to WT. Data represent mean ± s.d. of three independent biological replicates. Statistical analyses were performed using a ratio-paired t-test between WT and the individual mutant strains. p-value summary: \**P* < 0.05; \*\**P* < 0.01, \*\*\*\**P* < 0.0001; ns, not significant. **c**, Steady-state WT RhlR and ΔN17 RhlR levels at 3, 6, and 24-hours of growth. Samples were collected at the indicated times and analyzed by immunoblotting with anti-RhlR antibody. Band intensities were normalized to total protein staining and expressed relative to WT at each timepoint. Data represent mean ± s.d. of three independent biological replicates. Statistical analyses were performed using a ratio-paired t-test compared to WT RhlR levels at the indicated time points. p-value summary: \**P* < 0.05; \*\**P* < 0.01, \*\*\*\**P* < 0.0001; ns, not significant.

We next tested whether PqsE-dependent protection of RhlR influences RhlR steady-state levels. Overexpression of *pqsE* in a Δ*pqsE* background increased RhlR abundance by approximately ten-fold relative to the empty vector control (**Fig. 4b**), consistent with a stabilizing effect *in vivo*. Suggesting that PqsE plays a primary role in RhlR stability.

To determine whether degron-mediated turnover influenced RhlR abundance throughout growth, we monitored RhlR levels in wild-type and ΔN17 RhlR strains at 3, 6, and 24 hours following back-dilution (**Fig. 4c**). Strikingly, the strain expressing the ΔN17 variant accumulated significantly more RhlR than wild type at both 3 and 6 hours (**Fig. 4c**). However, this difference was lost after 24 hours, suggesting that the contribution of degron-mediated proteolysis changes across growth. Changes in expression of regulators that influence RhlR stability, including *pqsE* and *clpP2*, may contribute to these dynamics (**Supplementary Fig 5**). Because RhlR abundance is normally tightly restricted during early growth, the ΔN17 mutant provides a genetic tool to investigate how premature accumulation of RhlR influences QS-dependent transcription and downstream phenotypes.

### Stabilized RhlR accelerates activation of the RhlR regulon

To determine whether increased RhlR stability alters downstream QS activity, we examined activation of RhlR-dependent promoters. Because the ΔN17 variant accumulates elevated RhlR levels during early growth (**Fig. 4c**), we hypothesized that stabilization of RhlR would advance the onset of QS-dependent transcription. To test this, we first examined the activation of the phenazine genes, the biosynthetic genes whose products lead to the synthesis of pyocyanin which are directly regulated by RhlR^17^. We utilized chromosomal mScarlet transcriptional fusions of the *phzA1* and *phzA2* promoters and integrated them into wild-type and the strain expressing ΔN17 RhlR. We then monitored fluorescence following back-dilution into fresh medium. Consistent with our hypothesis, both reporters showed earlier and enhanced activation in the strain expressing the ΔN17 RhlR variant (**Fig. 5a**). Reporter activity began increasing between 3 and 4 hours post-back-dilution in cultures expressing ΔN17 RhlR variants, whereas wild-type cultures remained near baseline during this period. By 6 hours, expression from both promoters was significantly elevated in the strain expressing ΔN17 RhlR relative to wild type. In contrast, the differences dissipated by 24 hours, consistent with the convergence of RhlR protein levels observed at this time point (**Fig. 4c**). These findings indicate that degradation of RhlR and PqsE-mediated protection delays activation of QS-regulated promoters during early growth.

**Fig. 5:**
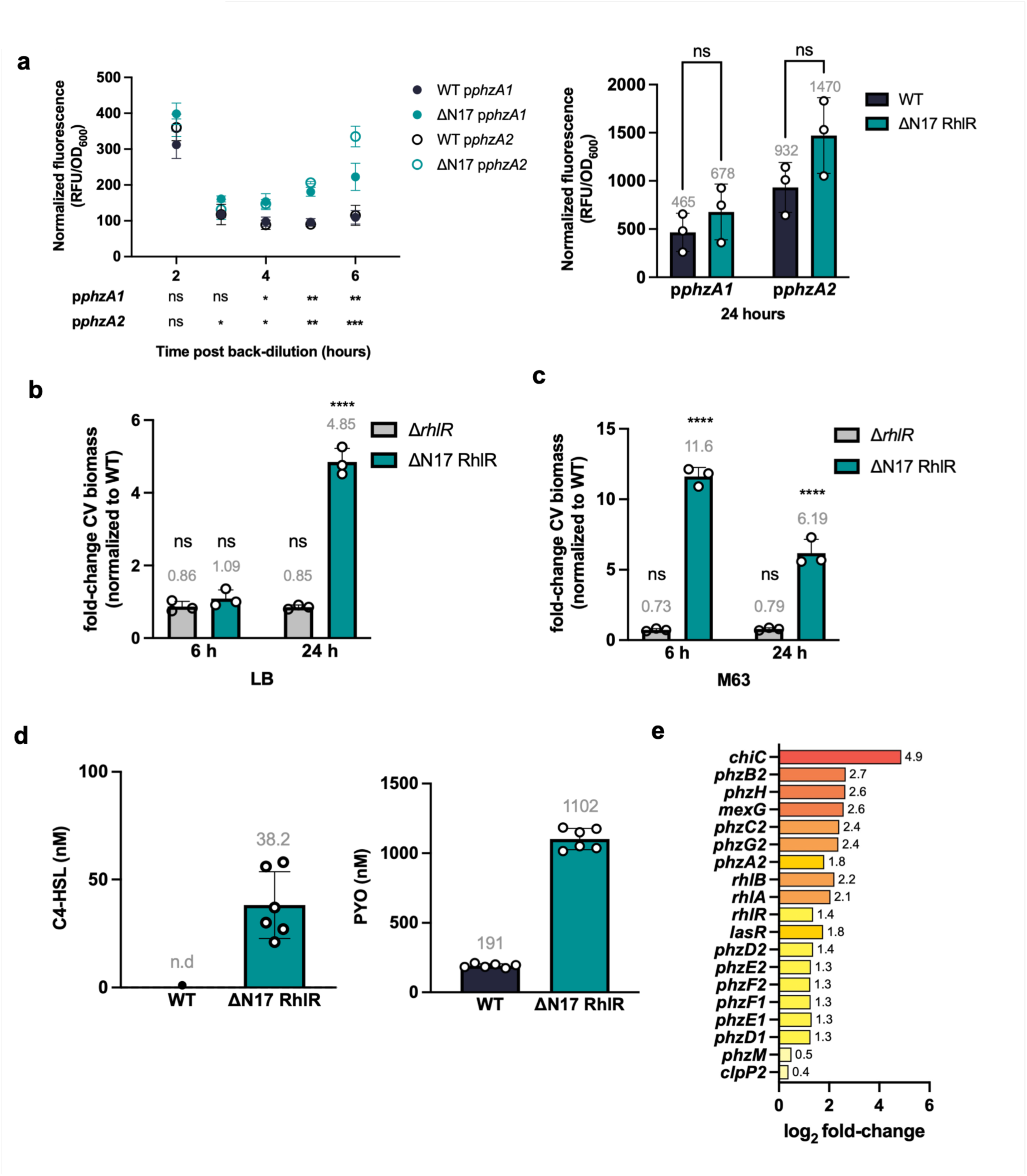
Stabilized RhlR enhances biofilm formation and secondary metabolite production. **a**, Activation of p*phzA1*-mScarlet and p*phzA2*-mScarlet in WT PA14 and ΔN17 *P. aeruginosa* strains is represented as fluorescence (ex, 520 nm; em, 580-640 nm) divided by OD600. The measurements are shown across a six-hour time course post back-dilution. Data represent three independent biological replicates with error bars showing the standard error. Statistical significance of ΔN17 vs. WT at each time point was determined by two-way ANOVA with Šídák’s multiple comparisons test (\**P* < 0.05, \*\**P* < 0.01, \*\*\**P* < 0.001). **b**, Crystal violet (CV) biofilm quantification in LB medium at 6 and 24 hours. Data are expressed as fold-change normalized to WT (set to 1) where CV staining was measured by OD560. Statistical significance was determined by one-way ANOVA with Dunnet’s multiple comparison test; \*\*\*\**P* < 0.0001; ns, not significant. **c**, Same as in panel b, with the exception that experiments were performed in M63 medium. **d**, C4-HSL and pyocyanin were quantified in culture supernatants after 3 h of growth by LC-MS using standard curve fitting. Data points represent 3 technical replicates from each of 2 biological replicates (6 points total per strain). Bars represent the mean; error bars represent SD. C4-HSL was not detected (n.d.) in WT supernatants (below the assay limit of detection). Given the limited number of biological replicates (n = 2), formal statistical comparisons were not performed. **e**, RNA-seq analysis showing log₂ fold-change in gene expression of ΔN17 relative to WT at 3 hours of growth. Genes are ranked by magnitude of fold-change. Highlighted in red are the most highly upregulated genes. At 6 and 24 hours, RhlR-regulated genes show minimal differential expression between ΔN17 and WT (**Dataset 1**). Data represent two independent biological replicates. All expression values are normalized to WT at the corresponding timepoint.

To determine whether accelerated transcriptional activation produces measurable physiological consequences, we next quantified pellicle biofilm formation via a crystal violet assay in wild-type, Δ*rhlR*, and ΔN17 RhlR variant strains. In LB medium, the ΔN17 RhlR variant and wild-type strains formed comparable biofilms after 6 hours of static growth, but by 24 hours ΔN17 RhlR variant strains produced significantly more biofilm biomass than wild type (**Fig. 5b**). The effect was more pronounced in M63 minimal medium, where biofilm formation is typically more robust^63^ (**Fig. 5c**). In this medium, the strain expressing ΔN17 RhlR showed significantly enhanced biofilm formation at both 6 and 24 hours (**Fig. 5c**). In contrast, the Δ*rhlR* strain produced biofilm levels similar to wild type under all conditions tested, consistent with previous reports indicating that RhlR is not essential for static biofilm formation^64^. Together, these results suggest that elevated RhlR abundance enhances biofilm development without being strictly required for it. Consistent with increased RhlR activity, the ΔN17 RhlR variant strain also produced modestly elevated levels of rhamnolipids and increased swarming motility (**Supplementary Fig. 6a, b**).

To test if the transcriptional changes we observed in the *phz* reporters led to altered metabolite production, we quantified QS-associated metabolites by liquid chromatography-mass spectrometry throughout growth. Consistent with the reporter assays, ΔN17 RhlR variant cultures accumulated substantially higher levels of several RhlR-regulated metabolites during early growth (**Fig. 5d**). C4-HSL and PYO were both elevated in the ΔN17 RhlR variant strain at 3 hours relative to wild type, indicating that stabilization of RhlR accelerates production of downstream QS outputs. Notably, several of the differences between the two strains observed at earlier time points were reduced or reversed after 24 hours of growth, including C4-HSL and PYO production (**Supplementary Fig. 6c, d**). This pattern is consistent with the convergence of RhlR protein levels and reporter activity observed at later time points. Thus, stabilization of RhlR primarily affects the timing of metabolite production rather than the ultimate accumulation of these molecules.

To determine whether the effects observed for individual promoters and metabolites reflected broader changes across the RhlR regulon, we performed RNA-seq on wild type and the strain expressing ΔN17 RhlR harvested at 3, 6, and 24 hours following back-dilution. At 3 hours, numerous established RhlR-dependent genes were upregulated in the ΔN17 RhlR variant strain, including the *rhl* cluster (*rhlA* and *rhlB*) and phenazine biosynthesis genes, which showed up to a 6-fold increase in expression (**Fig. 5e**; **Table 1**). In contrast, little to no differences were observed at the later time points, indicating that stabilization of RhlR primarily alters the timing of QS activation rather than the final magnitude of the transcriptional response (**Dataset 1**). Together, the reporter assays, metabolomic analyses, physiological phenotypes, and transcriptomic data demonstrate that proteolytic regulation of RhlR delays activation of the RhlR regulon during early growth and is required for its timely transition between low-cell-density and high-cell-density behaviors.

### Failure to degrade RhlR impairs competitive fitness

Because stabilized RhlR accelerated QS activation across multiple outputs, we next asked whether preventing RhlR degradation influences competitive fitness. To determine if premature activation of QS influences bacterial fitness, we performed mixed-culture competition assays between wild-type and ΔN17 RhlR variant strains under shaking and static growth conditions in both LB and M63 medium. Wild-type cells expressing constitutive mScarlet were mixed 1:1 with ΔN17 RhlR variant cells expressing constitutive mNeonGreen and plated immediately to determine the initial population ratio. Cultures were then grown for 24 hours prior to CFU enumeration of each fluorescent population. Across all tested conditions, the ΔN17 RhlR variant strain exhibited a reproducible fitness disadvantage relative to wild type (**Fig. 6a**). During planktonic growth in LB and M63 medium, the ΔN17 RhlR variant strain displayed mean log2 competitive indices (log2CI) of -1.70 and -1.74, respectively, corresponding to approximately three-fold fitness defects. Under static growth conditions, the ΔN17 RhlR variant strain exhibited a more modest defect in LB (mean log2CI = -1.08; ∼2-fold), with the strongest defect in M63 medium (mean log2CI = -2.80; ∼7-fold). Thus, stabilization of RhlR imposes a fitness cost across diverse environmental conditions. To ensure that these results were not influenced by the fluorescent markers, we repeated the competitions with the fluorophores reversed (WT-mNeonGreen versus ΔN17-mScarlet). Similar fitness defects were observed in all conditions tested (**Supplementary Fig. 7a**), demonstrating that the competitive disadvantage arises from the ΔN17 mutation itself rather than marker-dependent effects.

**Fig. 6:**
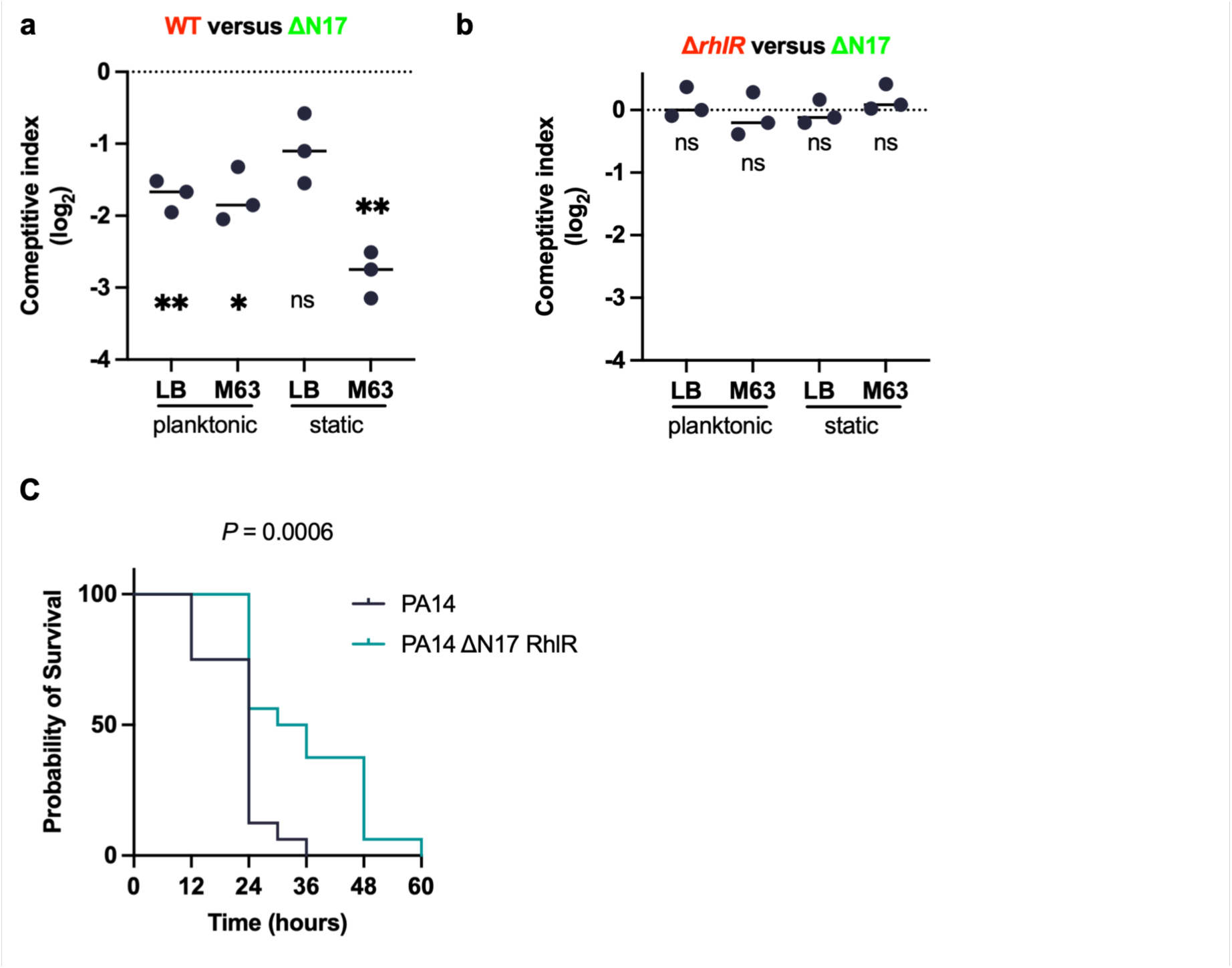
Stabilized RhlR imposes a competitive fitness cost *in vitro* and reduces *P. aeruginosa* virulence in a murine acute infection model. a, Competitive fitness assay comparing WT PA14 and ΔN17 RhlR strains across various growth conditions. WT cells constitutively expressing mScarlet were mixed 1:1 with ΔN17 RhlR cells expressing mNeonGreen, grown for 24 hours under planktonic (LB and M63 media) or static (LB and M63 media) conditions, and enumerated by CFU plating. Data are represented as log₂ competitive index (CI), where negative values indicate fitness defects in ΔN17 relative to WT. Each point represents an independent biological replicate. Statistical significance was determined by one-sample t-test and Wilcoxon test comparing log₂-CI values to 0 (no fitness defect); **; *P* < 0.01; \*\*\**P* < 0.001. Reciprocal labeling experiments with swapped fluorophores (**Supplementary** Fig. 8) confirm these results are independent of fluorescent markers. **b**, Same as in a, except the competition was between Δ*rhlR* and ΔN17 RhlR. **c**, Survival of C57BL/6 mice after intra-tracheal infection with 1×10^6^ CFU of WT PA14 or PA14 ΔN17 RhlR. Mice were monitored every 12 h for mortality. Statistical significance analyzed by Mantel-Cox test; n=16 mice per group; \*\*\**P* < 0.001.

The broad nature of the fitness defect suggested that stabilization of RhlR imposes a cost across diverse growth conditions. Because the ΔN17 RhlR variant strain exhibits elevated expression of multiple QS genes and metabolites, we hypothesized that wild-type cells benefit from exploiting public goods produced by the stabilized strain. To test whether the competitive advantage required the ability to respond to QS outputs, we repeated the competition assays using a Δ*rhlR* strain in place of wild type. In contrast to competitions with wild type, the ΔN17 RhlR variant and Δ*rhlR* strains showed no difference in competitive fitness across all conditions tested (**Fig. 6a, Supplementary Fig. 7b**). Thus, the competitive disadvantage associated with stabilized RhlR depends on the presence of a functional RhlR signaling system in the competing population (**Fig. 6b**).

Furthermore, we wanted to test the difference in pathogenicity of wild-type and strains expressing ΔN17 RhlR. To do this, two human bronchial epithelial cell lines, 16HBE14o- and CFBE41o- (normal and CF models respectively), were infected with wild-type and the ΔN17 RhlR variant at an MOI of 10. When measuring the cytokine production of these infections, there was no biologically significant differences in the secretion of IL-8, IL-6, nor TNF between wild-type and the strain expressing ΔN17 RhlR in either model (**Supplementary Fig 8**). These data indicate that while both the wild-type RhlR and stabilized RhlR are pro-inflammatory when compared to the uninfected control, they show very little difference between one another in virulence potential in a cell culture model system of the bronchial epithelium.

To determine the effect of ΔN17 RhlR on bacterial virulence, C57BL/6 mice were intra-tracheally infected with wild-type PA14 or PA14 expressing ΔN17 RhlR at an infectious dose of 1×10^6^ per mouse and survival of mice was monitored every 12h. While there was 100% lethality observed in both groups, mice infected with the wild-type PA14 strain succumbed to infection significantly faster withwn 36h after infection compared to mice infected with the PA14 ΔN17 RhlR strain that survived until 60h after infection (**Fig. 6c**) . This indicates that the ΔN17 RhlR strain was less virulent than the wild-type strain.

## DISCUSSION

Here, we identify regulated proteolysis as a previously unrecognized layer of QS control in *P. aeruginosa.* We demonstrate that the LuxR-type transcription factor RhlR is subject to regulated proteolytic turnover mediated by Lon and the ClpP1-ClpP2 complex through an N-terminal degron that is protected from protease recognition by its co-regulator PqsE. Preventing this turnover by removing the RhlR degron (ΔN17 RhlR) increased RhlR abundance, advanced the timing of QS activation, imposed a measurable competitive fitness cost, and reduced virulence in an acute mouse lung infection. Together, these findings establish that RhlR abundance is actively controlled by proteolysis and reveal receptor turnover as an important, regulated determinant of QS regulation.

Our data establish that RhlR is degraded by at least two ATP-dependent proteolytic systems, Lon and the ClpP1-ClpP2 heterocomplex (**Fig. 1**). A notable finding is the paradoxical stabilization of RhlR in *clpP2* single-deletion strains despite ClpP2 lacking independent proteolytic activity. This observation is explained by the biochemistry of the *P. aeruginosa* Clp system. As ClpP2 accumulates during stationary phase^47,65^, a substantial fraction of ClpP1 transitions from its homo-oligomeric form into the ClpP1-ClpP2 heterocomplex which exhibits distinct substrate specificity^46^. Complementation of *clpP1* alone in the Δ*clpP1* Δ*clpP2* background partially restored RhlR turnover, whereas *clpP2* alone could not, demonstrating that ClpP1 provides the catalytic activity, while ClpP2 confers the substrate specificity required for RhlR degradation (**Fig. 1d**). Although both Lon and ClpP1-ClpP2 can recognize RhlR, we found no evidence for functional compensation between these proteolytic systems under native expression conditions. Instead, restoration of RhlR degradation in the Δ*clpP1* Δ*clpP2* mutant required overexpression of *lon*, indicating that elevated Lon levels can substitute for the loss of *clpP1* and *clpP2,* but that endogenous Lon is insufficient to do so (**Fig. 1d**). These findings argue against redundancy and instead suggest that the two proteases contribute to RhlR turnover under distinct physiological or environmental conditions. Defining the cellular cues that bias RhlR towards one proteolytic pathway or the other will be important for understanding how proteolysis is integrated with QS regulation. We have not ruled out the contributions from other ATP-dependent proteases in *P. aeruginosa* (e.g., HslUV, FtsH, and the recently described Lon homolog AsrA^66^). Additionally, the unfoldase partner(s) that couple with ClpP1/ClpP2 for RhlR degradation remain to be defined.

A central finding of this work is that RhlR abundance is actively constrained by a self-limiting negative feedback loop (**Fig. 2**). RhlR directly activates transcription of *clpP2*, encoding a subunit of the protease complex responsible for its own degradation, while simultaneously repressing *pqsE*, which encodes the factor that protects RhlR from protease recognition (**Fig. 2a**). The RhlR K217A/K221A DNA-binding mutant confirms this architecture, exhibiting reduced *clpP2* expression, elevated *pqsE* expression, and increased RhlR stability (**Fig. 2d, 2e**). Thus, RhlR activity establishes opposing regulatory outputs that limit its own abundance. Consistent with this architecture, *clpP2* transcript levels were elevated in the strain expressing ΔN17 RhlR, as expected if stabilized RhlR activates expression of its own protease subunit. Because ΔN17 RhlR receptor lacks the degron, the additional ClpP1/P2 cannot act on RhlR, and the negative feedback loop is effectively uncoupled. These findings suggest that RhlR proteolysis functions to not only decrease receptor abundance but to alter the temporal dynamics of QS activation.

Identification of the N-terminal degron provides a mechanistic explanation for how PqsE stabilizes RhlR *in vivo* (**Fig. 3**). Structural analysis places residues 1-17 at a solvent-exposed region of the PqsE dimer interface, where a cluster of hydrophobic residues is well positioned to serve as a recognition element for ATP-dependent proteases. Removal of this region completely stabilized RhlR while preserving RhlR-PqsE-dependent pyocyanin production, indicating that degron removal does not disrupt productive interaction with PqsE. The stabilization of ΔN17 RhlR was independent of the presence of PqsE. We therefore propose that PqsE functions to physically occlude the protease recognition element. This mechanism is conceptually analogous to ligand-mediated stabilization of canonical LuxR-family receptors, but it differs in that protection is provided through a protein-protein interaction rather than ligand occupancy. This model also reconciles why RhlR was absent from previous proteomic surveys of Lon and ClpXP substrates^40,43^. Those studies were performed during early logarithmic growth, when PqsE-mediated shielding of the degron would be expected to limit protease access, suggesting that failure to detect RhlR reflects the physiological conditions examined rather than an absence of proteolytic regulation. Whether comparable degrons exist in other LuxR-type receptors remains to be determined. Protease susceptible may be common among unliganded LuxR-type receptors, TraR is degraded by Lon and Clp proteases in the absence of AHL^38^, LuxR accumulates in *E. coli lon* mutants expressing the *V. fischeri lux* system^67^, and PpuR levels rise in a *P. putida lon* mutant^68^. In *P. aeruginosa*, Lon affects the LasR/LasI system, but this appears to act through LasI, and direct proteolysis of LasR has not been shown^42^. In contrast, protection by a PqsE-like protein partner and specificity for ClpP1/ClpP2 may be particular to RhlR.

Our steady-state analyses demonstrated that degron-mediated proteolysis actively constrains RhlR abundance during early growth and that this constraint is diminished as cultures mature (**Fig. 4**). The convergence of wild-type and ΔN17 RhlR protein levels by 24 hours indicates that proteolysis primarily influences the transition into high-cell-density behaviors rather than the final magnitude of QS activation. The convergence of RhlR levels may reflect growth-stage-dependent changes in RhlR production or global protease activity that ultimately compensate for differences in degron-mediated degradation. Consistent with this model, stabilization of RhlR advanced activation of transcriptional reporters, increased production of QS metabolites, enhanced biofilm formation, and altered global transcription predominantly during early growth with these differences largely disappearing at later time points (**Fig. 5**). These observations indicate that proteolytic regulation governs the timing of QS activation rather than its eventual endpoint. One mechanistic explanation for this phenotype is the increased signaling sensitivity conferred by elevated receptor abundance. Previous work demonstrated that overexpression of RhlR lowers the apparent EC50 for C4-HSL^69^, suggesting that receptor abundance itself influences the signaling threshold. By permitting premature accumulation of RhlR, the ΔN17 mutation likely lowers the concentration of C4-HSL required to activate QS genes, thereby advancing the onset of the QS response.

The competitive fitness experiments provide an evolutionary explanation for why RhlR degradation is a highly regulated process (**Fig. 6**). Stabilization of RhlR imposed a reproducible fitness cost across both shaking and static growth conditions, and this disadvantage disappeared when the competing population lacked a functional RhlR signaling system. These findings are consistent with exploitation of prematurely produced QS public goods by neighboring cells and extend previous observations that hyper-QS mutants are disadvantaged in mixed populations^70^. Thus, regulated receptor turnover appears to balance the benefits of rapid QS activation against the energetic and ecological costs associated with activating cooperative behaviors too early. One caveat is that cells expressing ΔN17 RhlR express elevated *clpP2*. Increased protease activity could alter the turnover of other ClpP1/ClpP2 substrates and thereby contribute to some ΔN17 phenotypes independently of RhlR abundance. We cannot exclude this possibility, although the loss of the fitness cost when competitors lack functional RhlR signaling (**Fig. 6b**) indicates that the dominant effects arise from the RhlR regulon.

More broadly, our findings suggest that proteolytic turnover may contribute to the transition between high- and low-cell-density states. Current models of QS deactivation have largely emphasized passive dilution of extracellular autoinducers, yet signal decay alone cannot readily account for rapid remodeling of transcriptional programs in fluctuating environments. Previous work has shown that the C4-HSL synthase RhlI is targeted for ClpXP-dependent degradation^42,45^, indicating that both synthesis of the QS signal and responsiveness to that signal are subject to post-translational control. Together with the regulated degradation of RhlR described here, these findings raise the possibility that ATP-dependent proteolysis acts at multiple nodes of the Rhl quorum-sensing circuit to actively tune signaling dynamics and facilitate circuit resetting.

To assess the relevance of this regulation during infection, we tested survival of PA14 and PA14 expressing ΔN17 RhlR in a C57BL/6 mouse intra-tracheal infection model. We observed significantly delayed mortality in the strain expressing ΔN17 RhlR compared to wild-type (**Fig. 6c**). The reduced virulence of the ΔN17 RhlR mutant *in vivo* suggests that QS timing plays an important role in the host pathogen interaction. Coupled with the discovery that the mutant strain is less competitively fit *in vitro*, there are several possibilities that could explain the delayed virulence of the mutant. It is possible that premature activation of the QS system and virulence factor production at low bacterial densities such as in the case of the ΔN17 RhlR mutant strain could increase activation of the immune system early on in infection and improve host immune cell clearance of the bacteria. Sub-lytic concentrations of rhamnolipids can stimulate immune cells^71^ and synthetic rhamnolipids are being explored as potential adjuvants^72^. A second possibility is that premature QS activation could be energetically costly to *P. aeruginosa* at low cell densities, causing the bacteria to grow more slowly and less able to compete with the host immune cells. However, this second possibility of a growth defect seems somewhat less likely since the ΔN17 RhlR mutant had no fitness cost relative to the Δ*rhlR* mutant, showing that the true competitive cost was due to cheating by wild-type *P. aeruginosa*. Regardless, these results indicate the receptor turnover is highly regulated (**Fig. 7**) further support targeting QS as a means to reduce bacterial virulence, showing that even disrupting timing of QS could lead to improved survival.

**Fig. 7:**
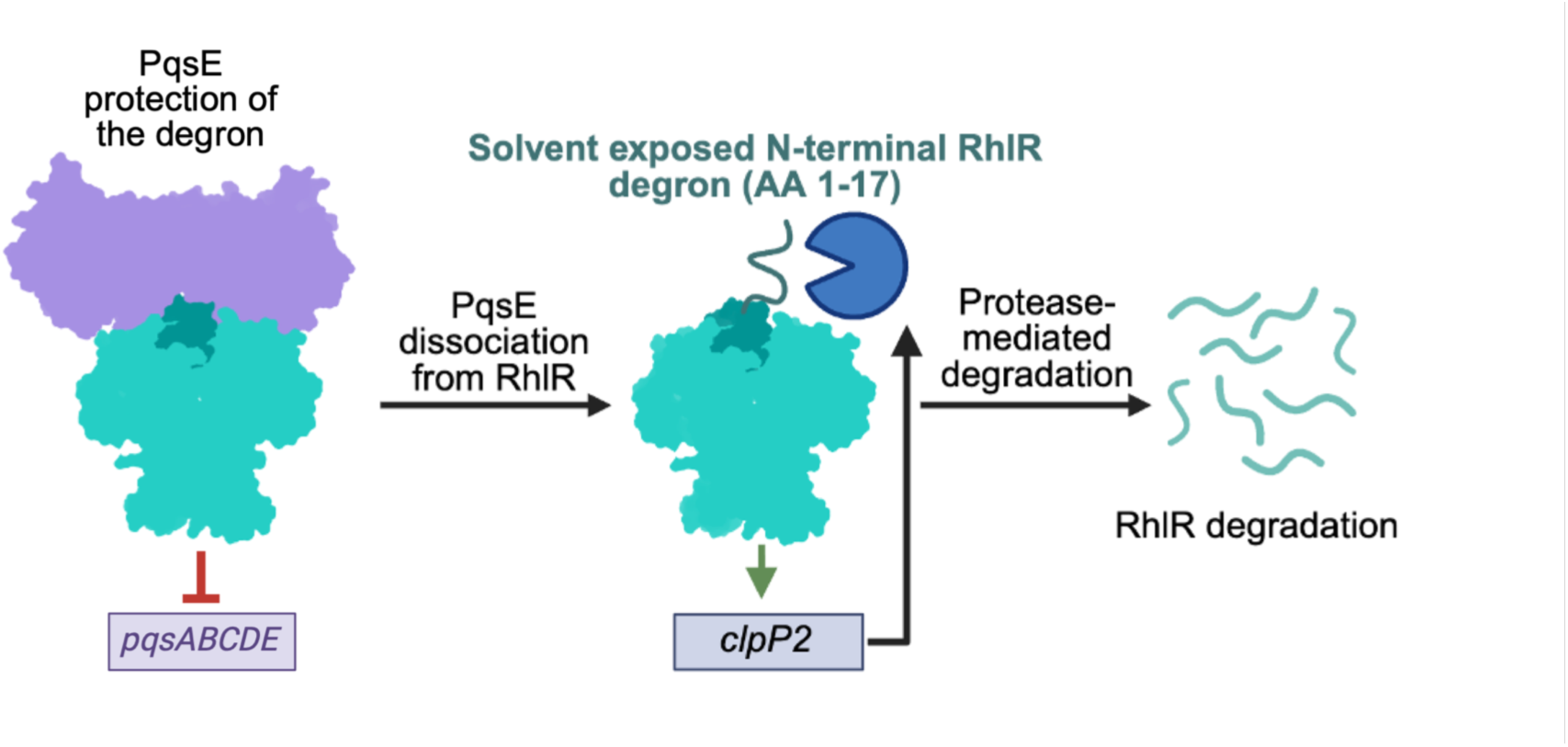
Model for PqsE-mediated protection and regulated proteolysis of RhlR. When PqsE is bound to RhlR, it shields the solvent-exposed N-terminal degron (amino acids 1-17) or RhlR from protease recognition (left). RhlR represses transcription of the *pqsABCDE* operon (red blunt, ended line), which encodes PqsE, limiting the abundance of its own protector. Upon PqsE dissociation from RhlR, the degron becomes exposed and accessible to ATP-dependent proteases (middle). RhlR also activates transcription of *clpP2* (green arrow), which encodes a subunit of the ClpP1/ClpP2 protease complex that degrades RhlR. Protease-mediated degradation of RhlR (right) completes a negative feedback loop that limits RhlR abundance.

## MATERIALS AND METHODS

### Strain construction

*P. aeruginosa* mutant strains were generated by allelic exchange using pEXG2 derivative plasmids, as previously described^73^. Chromosomal deletions of *clpP1*, *clpP2*, *clpX*, *clpA*, *clpS*, and the RhlR N-terminal 17 amino acid deletion, ΔN17 RhlR, were constructed following the pEXG2 strategy with plasmids independently introduced into PA14 via conjugal mating with *E. coli* SM10 λpir as the donor strain. The *rhlR* K217A/K221A strain was introduced into the PA14 chromosome using site-directed mutagenesis via pEXG2, following the same conjugation approach. Standard cloning and molecular biology techniques were used for plasmid construction in *E. coli* and *P. aeruginosa*.

### Colony forming unit enumeration

*P. aeruginosa* cells were cultured from glycerol stocks in 5 mL of LB at 37°C with shaking at 200 rpm. Overnight cultures were back diluted to an OD600 of 0.1 in 20 mL of LB and grown for 3, 6, and 24 h in 125 mL volumetric flasks. At each timepoint, 10 µL of *P. aeruginosa* cells were diluted 1:10 in a clear 96-well plate (CellTreat) in a total volume of 100 µL containing fresh LB medium and 10-fold serially diluted a total of eight times. For each dilution, 10 µL was plated on an LB plate and grown overnight at 37°C.

### Pyocyanin production assays

At each timepoint used for CFU enumeration (3, 6, and 24 h), 1.5 mL of *P. aeruginosa* culture was aliquoted into a microcentrifuge tube and pelleted at 21,300 × g for 2 min to pellet the cells. 1 mL of cell-free supernatants was transferred to a clear cuvette, and pyocyanin production was quantified by measuring the absorbance at 695 nm using an Ultrospec 3100 pro spectrophotometer (Amersham Biosciences). Absorbance values from sterile LB medium were subtracted as a background control.

### *In vivo* protein stability and Western blot analyses

Cells were cultured from glycerol stocks in 5 mL of LB at 37°C with shaking at 200 rpm. Overnight cultures were back diluted to an OD600 of 0.1 in 20 mL of LB and grown for 6 h or to an OD600 of ∼2.5 at 37°C with 200 rpm. For translation inhibition experiments, after 6 h, protein synthesis was inhibited by the addition of chloramphenicol (30 µg/mL final concentration). The cultures continued to incubate at 37°C with shaking at 200 rpm, in which 1.5 mL of cultures were collected at indicated timepoints (0, 30, 60, 90, and 120 min). Cells were harvested by centrifugation at 21,300 × g for 2 min and cell pellets were immediately resuspended in 2x SDS sample dye (250 mM Tris-HCl, pH 6.8, 8% w/v SDS, 40% glycerol, 8% β-mercaptoethanol, 0.02% bromophenol blue) normalized by OD600 values (50 µL of 2x SDS sample buffer / OD600 = 0.2) and flash frozen in liquid nitrogen before storage at -20°C. Lysed samples were boiled for 10 min at 95°C or 5 min at 100°C and centrifuged at 21,300 × g for 2 min. After centrifugation, 10 µL of the supernatant was run on Criterion TGX precast SDS-PAGE gels (Bio-Rad) at 35 mA for 35 min. Protein was transferred to polyvinylidene difluoride (Bio-Rad) membranes at 110 A for 1 h using a semi-dry transfer cell (Bio-Rad). Following protein transfer, blocking was performed with 1x Tris-buffered saline with Tween 20 (TBST) and 5% milk for 2 h at room temperature with agitation. Primary α-RhlR polyclonal antibody from rabbit (Cambridge Antibodies) was incubated with the membrane at a 1:1,000 dilution overnight at 4°C with agitation. Following incubation with the primary antibody, three 5-min washes were performed with TBST before incubation with goat anti-rabbit cross-adsorbed secondary antibody conjugated with horseradish peroxidase (Thermo Fisher Scientific) for 1 h at room temperature with agitation at a 1:10,000 dilution. Three 5-min washes with TBST were performed before the addition of Pierce ECL western blotting substrate (Thermo Fisher Scientific) and imaging with a 2-min exposure on an iBright-1500 (Thermo Fisher Scientific). Final images were quantified using ImageJ software and degradation rates were plotted using Prism 10.2.3 software. All antibody solutions were made in 1x TBST and 5% milk. Steady-state protein levels were measured exactly as above, with the exception that cells were not treated with chloramphenicol.

### Luciferase reporter assays

Luciferase reporter assays were performed as previously described. Briefly, *E. coli* strains harboring *rhlR* or *lasR* expressed from the pBAD promoter, together with *luxCDABE* under the control of the p*clpP2* or p*lasB* promoters in the pCS26 vector, and where indicated, either the pACYC184 vector alone or harboring *pqsE*, with expression from the *lac* promoter, were cultured from glycerol stocks in 5 mL of LB at 37°C with shaking at 200 rpm. Overnight cultures were back diluted 1:100 in 20 mL of LB and grown for 2 h or until OD600 ∼0.4-0.6. At this time, the cultures were induced with a final concentration of 0.1% arabinose and 198 µL of induced culture was added to the wells of a clear bottom black opaque 96-well plate (Corning) containing 2 µL of C4-HSL (RhlR) or 3OC12-HSL (LasR) or no HSL (arabinose only control). The final concentration of HSL in the wells was 10 µM. The 96-well plate was incubated at 37°C for 4 h with shaking before luminescence was measured using a Glomax (Promega) plate reader.

### Crystal violet assays

Overnight cultures were back-diluted 1:100 into fresh LB or M63 medium supplemented with 1 mM MgSO42-, 0.2% glucose, and 0.5% casamino acids into a clear 96-well plate (CellTreat) in at least six technical replicates. The plate was incubated at 37°C for 6 or 24 h. Cells were removed from the plate by inversion, and the wells were washed with sterile water to remove excess cells and media. After the washes, 125 µL of 0.1% crystal violet in water was added to each well and incubated at room temperature for fifteen minutes. After incubation, crystal violet was removed and the plate was washed thoroughly with sterile water and left to dry for several hours or overnight. After drying, 125 µL of 30% acetic acid was added to each well and incubated for 15 minutes. After solubilization, the crystal violet was transferred to a new 96-well plate and absorbance was read at 560 nM on a Glomax (Promega) plate reader with 30% acetic acid used as a blank control. For each biological replicate, technical replicates were averaged per strain and plotted as the fold-change relative to wild type.

### Swarming assays

Swarming plates were made with 5X M9 Salts (20 mM NH4Cl, 12 mM Na2HPO4, 22 mM KH2PO4, 8.6 mM NaCl), 0.4% agar, 0.5% casamino acids, 1 mM CaCl2, 1 mM MgSO4, and 1.35% (w/v) dextrose. Overnight cultures were grown in 3 mL of LB and normalized to an absorbance of OD600 ∼1.0. 5 µl of normalized culture was spotted onto the center of fresh swarming media and left to dry. Plates were incubated face down at 37°C for 24 h and then at room temperature for another 12 hours. Images of plates were taken using Epson Perfection V850 Pro scanner at a resolution of 300 dpi. Swarming plates were analyzed using our previously published protocol^18^.

### Rhamnolipid production

The wells of clear 96-well plates (CellTreat) were stained with 50 µL of 0.1 mg/mL Victoria Pure Blue BO dye in isopropanol. The isopropanol was evaporated under vacuum for 1 h at 45°C. Following evaporation, 300 µL of 0.5 M NaOH was added to the wells and incubated for 10 min at room temperature before drying under vacuum for 1 h at 45°C. The strains were grown overnight from single colonies overnight in 3 mL of LB medium at 37°C with shaking at 200 rpm. 1 mL of overnight cultures were aliquoted into microcentrifuge tubes and centrifuged for 2 min at 21,300 x g. Cell-free culture supernatants were added in technical triplicate to stained wells and incubated with agitation at 750 rpm for 1 hour at room temperature. After incubation, 200 µL was transferred to a new 96-well plate (CellTreat) and the absorbance was read at 625 nM (Molecular Devices SpectraMax M5 SoftMax Pro5).

### mScarlet transcriptional reporter assays

Strains containing mScarlet *phz1* or *phz2* reporters were grown overnight in 3 mL of 1% tryptone broth. Overnight cultures were back-diluted in 20 mL of 1% tryptone broth to an OD600 ∼ 0.1 in a 125 mL flask. Cultures were grown at 37°C with shaking at 200 rpm, after 2 h of growth 200 µL of cells were added to a clear bottom black opaque 96-well plate (Corning) where mScarlet fluorescence and optical density at 600 nm was measured using a Glomax (Promega) plate reader with excitation at 520 nm and emission collected across 580-640 nm for 2-6 h, and 24 h post back-dilution.

### RNA extraction and sequencing

#### RNA extraction

WT and ΔN17 strains were grown overnight from single colonies in 3 mL of LB medium at 37°C with shaking at 200 rpm. After overnight growth, cultures were back-diluted in 20 mL to an OD600 ∼ 0.1 in a 125 mL flask. After 3, 6, or 24 h of growth, 1.5 mL of cells were harvested by centrifugation at 21,300 x g for 2 min and pellets were frozen at -80°C until RNA extraction. RNA extraction was performed using the Invitrogen PureLink RNA Mini Kit. Cell pellets were thawed and resuspended in 100 µl of lysozyme solution containing 10 mM Tris-HCL pH 8.0, 0.1 mM EDTA, and 1 mg/mL lysozyme. 5 µl of 1% SDS was added, mixed by vortex and incubated at room temperature for 5 min. After the incubation, 350 µl of lysis buffer (Invitrogen) supplemented with 2-mercaptoethanol (10 µl of 2-mercaptoethanol per 1 mL of lysis buffer) was added to each sample, mixed by vortex, and transferred to screw-top tubes. Cells were lysed by bead beating (BioSpec 0.1 MM Zirconia/Silica Beads) for two rounds of 50 s and centrifuged at 12,000 x g for 2 min. The supernatant was aspirated into a fresh 1.5 mL tube containing 250 µl of 100% ethanol and mixed by vortex. Samples were transferred to a spin column (Invitrogen), centrifuged at 12,000 x g for 20 s. Following centrifugation, the flow-through was discarded and the column was washed with 700 µl of Wash Buffer I (Invitrogen) by centrifugation at 12,000 x g for 20 s and washed twice with 500 µl of Wash Buffer II (Invitrogen) by centrifugation at 12,000 x g for 20 s. After the final wash, the flow-through was discarded and the samples were centrifuged at 12,000 x g for 1 min to dry the membrane bound RNA. A total of 31 µl of RNAse-free water (Invitrogen) was added to the spin column, incubated at room temperature for 1 minutes, and centrifuged at 12,000 x g for 1 min. Eluted RNA was frozen at -80°C until use.

#### RNA sequencing and data analysis

Bulk RNA was sent to SeqCenter (Pittsburgh, PA) on dry-ice where it was processed for paired-end Illumina sequencing by rRNA depletion using Ribo-Zero Plus (Illumina). Fastq files generated by SeqCenter were processed using Rockhopper v. 2.0.3 with default settings as recently described^18^.

### RT-qPCR

Total RNA from strains was prepared as described above and treated with the Turbo DNase kit per manufacturer’s instructions (ThermoFisher). cDNA was prepared with a SuperScript IV Reverse Transcriptase kit (Invitrogen) in a total reaction volume of 20 µl. Reactions containing 2X SYBR Select Master Mix (Applied Biosystems), primers (200 nM final concentration), and 2 µl of 1:5 diluted cDNA was added to a MicroAmp 96-well plate (Applied Biosystems) in a final volume of 20 µl per well. A 7500 Fast Real-Time PCR system (Applied Biosystems) was used for cycle threshold quantification. The *P. aeruginosa proC* gene was chosen as the internal housekeeping control to normalize the data and calculate the relative fold-change in gene expression using the 2^-ΔΔCt^ method.

### Direct ChIP-qPCR

Overnight cultures were back-diluted in 20 mL of LB to an OD600 ∼ 0.1 in a 125 mL flask and grown at 37°C with shaking at 200 rpm until the cultures reached an OD600 ∼ 2.0. Cells were crosslinked by the addition of 500 µl of formaldehyde for a final concentration of 1%. Cultures were swirled to mix and left to sit on the bench for 20 min at room temperature. Crosslinked cultures were quenched by transfer to 50 mL conical tubes containing 10 mL of 2.5 M glycine (0.5 M final concentration). Cells were pelleted at 21,300 x g for 5 min, the supernatant was discarded, and the cell pellets were resuspended in 20 mL 1x TBS and centrifuged for another 5 min at 21,300 x g. The supernatant was again discarded, and the cell pellets were resuspended in 1 mL of 1x TBS, transferred to microcentrifuge tubes for centrifugation at 21,300 × g for 5 min, and the cell pellets were flash frozen and stored at -80°C until further use. Thawed cells were resuspended in 1 mL of 150 mM FA lysis buffer (1 M HEPES, pH 7.5, 2 M NaCl, 0.5 M EDTA, 0.1% Triton-X 100, 5% sodium deoxycholate, 10% SDS, and sterile water) containing 4 mg/mL lysozyme (Sigma) and incubated at 37°C for 30 min. Cells were lysed by sonication at 0-10°C for 30 min with 30 s on/off pulses at 75% output using a BioRupter Sonicator (Diagenode). Cell lysates were centrifuged twice for 10 min at 21,300 x g, between each centrifugation, the cell-free supernatant was transferred to a new microcentrifuge tube. Following centrifugation, 1 mL of supernatant was diluted in 1 mL of 150 mM FA lysis buffer for a final volume of 2 mL. At this time, 100 µl of cross-linked, sonicated lysates were diluted in 800 µl of 150 mM FA lysis buffer, and 20 µl was saved as input samples. A total of 30 µl of a 50% slurry of protein A beads (Sigma) in 1 x TBS and 10 µl of primary α-RhlR polyclonal antibody from rabbit (Cambridge Antibodies) were added to lysates and incubated at room temperature for 90 min on a rotisserie rotator. Following incubation, the immunoprecipitation mix was centrifuged for 1 min at 6,000 x g, the supernatant was removed by aspiration, and the cell pellets were resuspended in 750 µl of 150 mM FA lysis buffer. This mix was transferred to a Spin-X column (Corning), incubated for 3 min at room temperature on a rotisserie rotator, centrifuged again for 1 min at 6,000 x g, and the supernatant was removed by aspiration. This was repeated with 700 µl of 150 mM FA lysis buffer, 500 mM FA lysis buffer, ChIP wash buffer (1 M Tris-HCl, pH 8.0, 1 M lithium chloride, 0.5 M EDTA, NP-40, and 5% sodium deoxycholate), and TE (10 mM Tris-HCl, pH 8.0 and 1 mM EDTA). After the final wash, protein A beads were resuspended in 100 µl ChIP elution buffer (1 M Tris-HCl, pH 7.5, 0.5 M EDTA, and 10% SDS). The Spin-X column was placed in a fresh dolphin-nosed tube (Sorenson) and incubated for 10 min at 65°C with occasional gentle mixing by inversion. Immunoprecipitation mixes were centrifuged for 1 min at 6,000 x g, and the Spin-X filters were discarded. Cells and input samples were de-crosslinked by incubation at 100°C for 10 min. DNA was cleaned up with a PCR purification kit (Qiagen). Input samples were diluted 1:100 for 10 µl qPCR reactions. qPCR reactions contained 5 µl SYBR Select Master Mix (ThermoFisher), 2 µl cleaned DNA, and 3 µl combined forward and reverse primers. qPCR reactions were run with a 50°C holding stage, a 95°C holding stage, 40 cycles of 95°C-60°C, and a continuous melt curve stage from 95°C to 60°C (Applied Biosystems 7500 Fast Real-Time PCR System).

### Liquid chromatography mass spectrometry of QS metabolites

#### Molecule extraction

Overnight cultures were back-diluted in 20 mL of LB to an OD600 ∼ 0.1 in a 125 mL flask. Strains were grown for 24 h with 1.5 mL aliquots taken at 3, 6, and 24 h for centrifugation at 21,300 x g for 2 min. Cell-free culture supernatants were aliquoted into new microcentrifuge tubes and frozen at -80°C until use. Sample supernatants were thawed and 200 µl was added to new microcentrifuge tubes containing 800 µl of methanol, vortexed for 15 s, and centrifuged for 2 min at 10,000 x g. Following centrifugation, the supernatant was mixed 1:1 with 0.1% formic acid in high-performance liquid chromatography vials and vortexed again.

#### Analytical measurements and analysis

C4-HSL and PYO were quantified by ultra-high-performance liquid chromatography coupled to high-resolution mass spectrometry (UHPLC-HRMS) using a method adapted from a previously described approach^18^. Briefly, samples were analyzed using a Vanquish UHPLC system coupled to an Orbitrap mass spectrometer (Thermo Fisher Scientific) operated in positive electrospray ionization mode. Chromatographic separation was performed using a Hypersil GOLD C18 column (50 × 2.1 mm, 1.9 µm) maintained at 45°C. Mobile phase A consisted of 0.2% formic acid in water, and mobile phase B consisted of 0.2% formic acid in acetonitrile. The flow rate was 0.45 mL/min. The 14-min gradient started at 5% B, increased to 100% B over 10 min, was held at 100% B for 1 min, and then returned to 5% B and re-equilibrated for 3 min. C4-HSL and PYO were identified by accurate mass and retention time relative to authentic standards. Peak areas were integrated, and C4 and PYO concentrations were determined using external calibration curves generated from authentic standards analyzed in the same batch.

### Competitive index assays

WT, ΔN17, and Δ*rhlR* strains constitutively expressing either mScarlet or mNeonGreen were grown overnight from single colonies in 3 mL of LB medium at 37°C with shaking at 200 rpm. After overnight growth, cultures were back-diluted and mixed 1:1 so that each culture contributed approximately OD600 0.05 in a final volume of 2 mL in a kaput tube for planktonic growth or 200 uL of a 96-well for static growth. After mixing, 10 µL of cells were diluted 1:10 in a clear 96-well plate (CellTreat) in a total volume of 100 µL containing fresh LB medium and 10-fold serially diluted a total of eight times. At time-point 0, dilutions ranging from 10^-4^ to 10^-6^ were plated on an LB plate and grown overnight at 37°C. The remaining competition mixture was incubated for 24 h at 37°C. After incubation 10^-7^ and 10^-8^ cells were plated on fresh LB medium and grown overnight at 37°C. Plates were imaged on a BioRad ChemiDoc MP Imaging System. mScarlet fluorescence was detected using Green Epi illumination with a 602/50 emission filter (Alexa 546 blot application setting) and a 3 s exposure time, and mNeonGreen fluorescence was detected using Blue Epi illumination with a 532/28 emission filter (Alexa 488 blot application setting) and a 1 s exposure time. The competitive index was calculated as the log2 ratio of the final number of red/green colonies/the ratio of the initial time 0 red/green colonies.

### Mammalian cell infection assay

16HBE14o- (Sigma Aldrich, #SCC150) and CFBE41o- (Sigma Aldrich, #SCC151) were maintained in EMEM supplemented 10% FBS, 1% L-glutamine, and 1% penicillin/streptomycin at 37 °C with 5% CO2. For cytokine detection, cells were seeded in 6-well dishes at 1×10^6^ cells per well and allowed to grow to confluency. 24 h prior to infection, cells were washed and EMEM supplemented with 10% FBS and 1% L-glutamine without antibiotic was added. Overnight cultures of *P. aeruginosa* were pelleted and washed three times in DPBS. Cells were normalized to an OD600 of 1.0 in DPBS. *P. aeruginosa* inoculated at an MOI of 10 and allowed to attach to the epithelium for 2 h. Cells were subsequently washed three times in DPBS to remove any unattached bacteria. Infection was allowed to proceed for an additional 4 h, at which time spent media was collected and stored at -80°C. CFU were enumerated to ensure similar growth across *P. aeruginosa* strains tested. The human inflammatory cytokine cytometric bead array (CBA) I kit (Waters, #551811) was used to determine the amount of IL-12p70, TNF, IL-10, IL-6, IL-1ß, and IL-8 present. Samples were prepared according to manufacturer’s instructions. Samples were read using a FACSymphony A3 Cell Analyzer (BD Biosciences).

### Animal infection studies

Overnight cultures of PA14 or PA14 ΔN17 RhlR were grown to mid-log phase in lysogeny broth and used for infections. 8-week old male or female C57BL/6 mice were obtained from the Jackson Laboratory and were housed in a specific-pathogen-free animal facility with 12:12 light: dark cycle, 21 ± 2 °C, ∼ 40% humidity at the Cedars-Sinai vivarium. For survival studies, mice were anesthetized using isoflurane and intra-tracheally infected with 1×10^6^ CFU of WT PA14 or PA14 ΔN17 RhlR. Infected mice were monitored every 12h for mortality and moribund mice were euthanized by carbon dioxide inhalation. All experiments were approved by the Institutional Animal Care and Use Committee at Cedars-Sinai Medical Center under protocol IACUC008115.

## DATA AVAILABILITY

Sequencing data have been deposited at the NCBI Sequence Read Archive under the BioProject accession number PRJNA1545364.

## SUPPORTING INFORMATION

**Figure S1.**
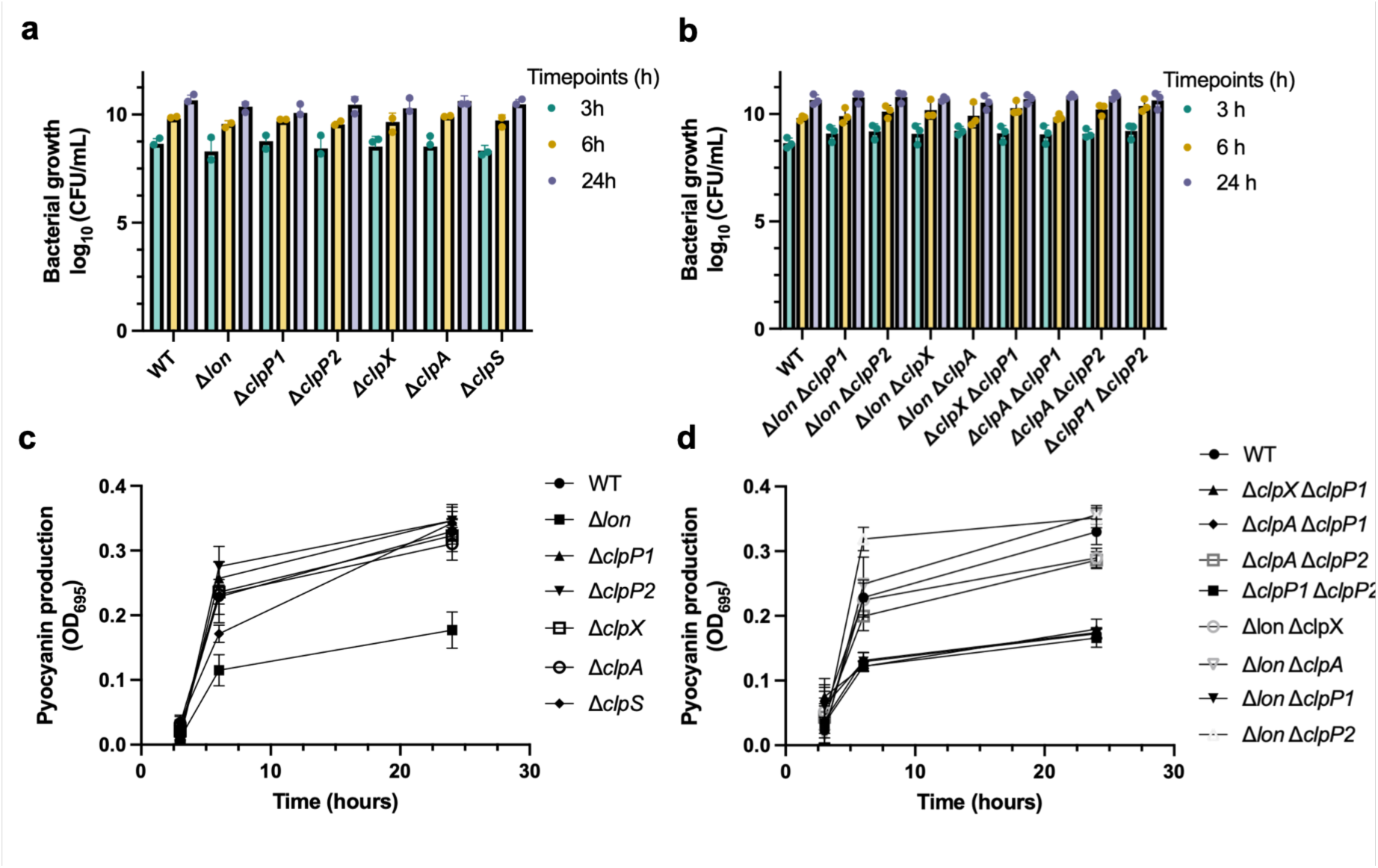
Growth and pyocyanin production are largely unaffected by protease deletions. a, Colony forming units (CFU/mL) of wild-type and indicated single (**a**) or double (**b**) protease mutants at 3, 6, and 24 hours of growth. **c,** Pyocyanin production measured by absorbance at OD695 of wild-type and indicated single (**c**) or double (**d**) protease mutants at 3, 6, and 24 hours. Data represents means ± SEM of n = 3 biological replicates. Statistical comparisons of each mutant to WT were performed by one-way ANOVA with Dunnett’s multiple comparisons test. Adjusted *P* values and 95% confidence intervals are provided in Supplementary Table 1.

**Figure S2.**
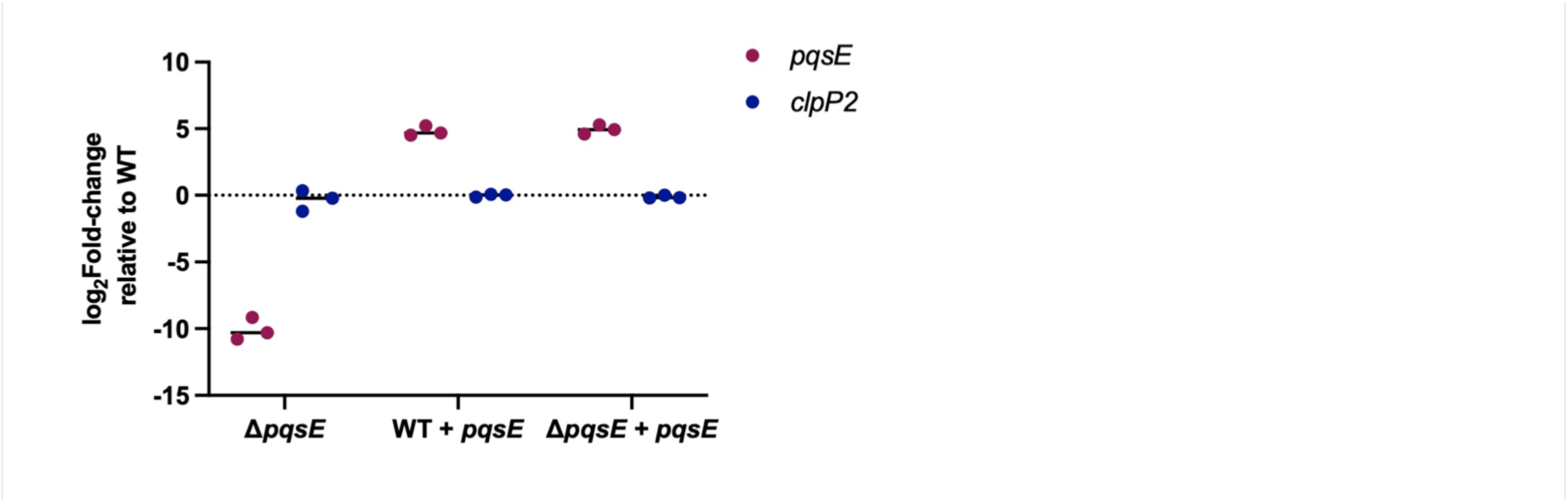
RT-qPCR analysis of *pqsE* and *clpP2* expression. Relative transcript levels of *pqsE* (magenta) and *clpP2* (blue) were quantified by RT-qPCR in the indicated strains grown to stationary phase (OD600 ∼2.0) and normalized to the housekeeping gene *proC* before comparison to wild-type. Deletion of *pqsE* abolished *pqsE* transcript detection, whereas plasmid-borne *pqsE* resulted in elevated *pqsE* expression in both wild-type and Δ*pqsE* backgrounds. *clpP2* transcript levels remained similar to wild type under all conditions tested, demonstrating that changes in RhlR stability associated with PqsE are not attributable to altered *clpP2* transcription. Each point represents one biological replicate.

**Figure S3.**
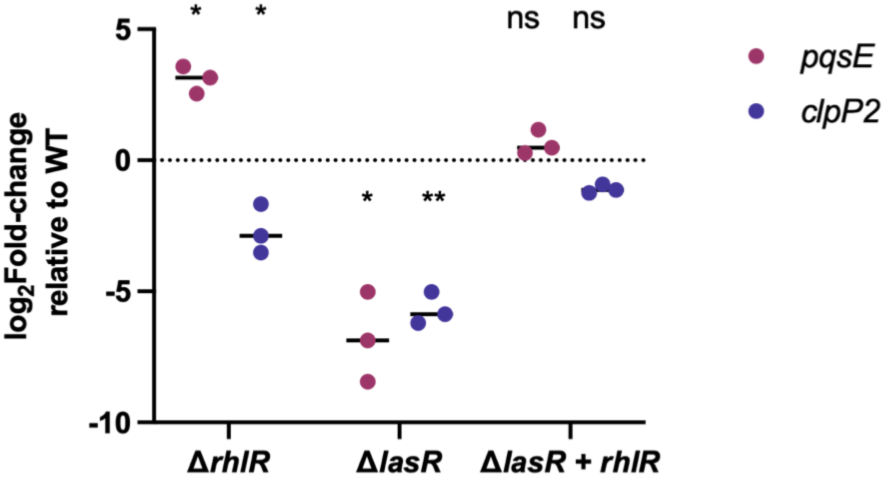
*lasR* and *rhlR* differentially regulate *pqsE* and *clpP2* transcription. RT-qPCR analysis of *pqsE* (magenta) and *clpP2* (blue) transcript levels in the indicated QS regulator mutants relative to wild-type in cells grown to stationary phase (OD600 ∼2.0). Deletion of *rhlR* increased *pqsE* transcript abundance while reducing *clpP2* expression, whereas deletion of *lasR* decreased expression of both genes. The Δ*lasR* Δ*rhlR* double mutant restored *pqsE* and *clpP2* transcript levels to near wild-type levels. Each point represents an independent biological replicate. Statistical significance was determined by one-way ANOVA with Dunnet’s multiple comparison test.

**Figure S4.**
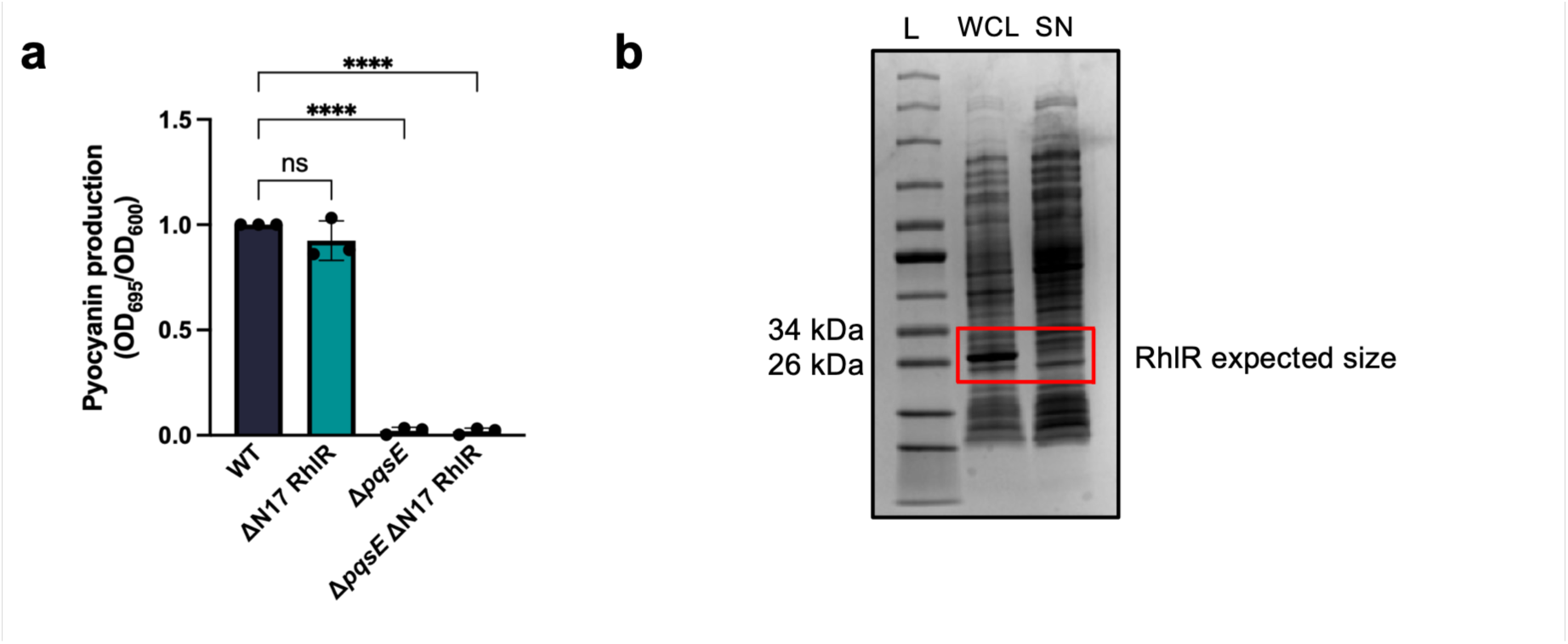
Additional controls for ΔN17 RhlR. a, Pyocyanin production (OD695) normalized by optical density (OD600), expressed relative to WT in the indicated strains. Each point represents independent biological replicates (n = 3), and bars show the mean ± SD. Each strain was compared to WT by one-way ANOVA with Dunnett’s multiple comparisons test (ns, not significant; ****, adjusted *P* < 0.0001). **b,** Coomassie-stained SDS-PAGE gel of whole-cell lysate (WCL) and supernatant (SN) fractions. No protein band corresponding to the expected molecular weight of RhlR (∼28 kDa) was detected in the SN fraction, indicating that RhlR is not soluble under the conditions tested. L, molecular weight ladder.

**Figure S5.**
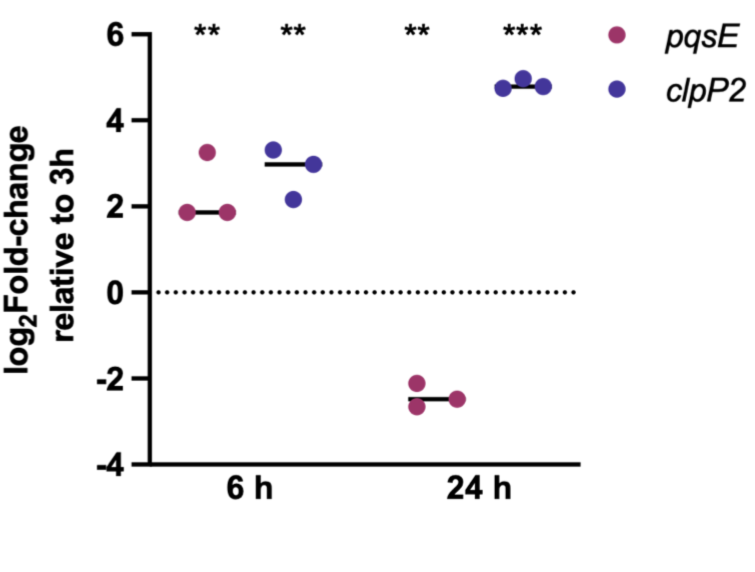
Temporal expression of *pqsE* and *clpP2* transcription. RT-qPCR analysis of *pqsE* (magenta) and *clpP2* (blue) transcript abundance at 6 and 24 hours relative to the 3-hour time point. *clpP2* expression increased throughout growth, whereas *pqsE* expression was elevated at 6 hours but decreased by 24 hours relative to 3 hours. Each point represents an independent biological replicate. Statistical significance was determined by one-way ANOVA with Dunnet’s multiple comparison test.

**Figure S6.**
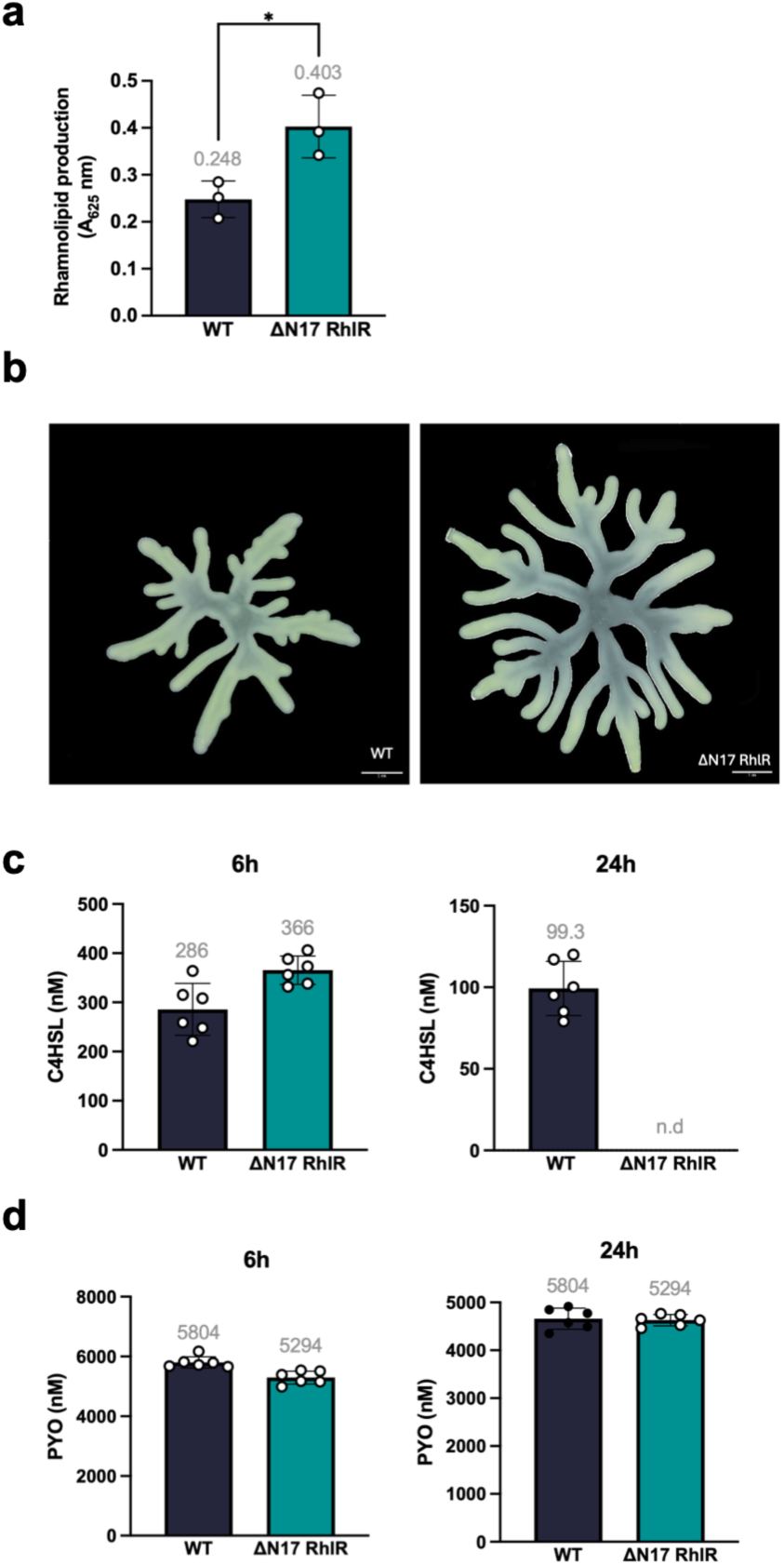
Rhamnolipid production and swarming motility are increased in the ΔN17 RhlR strain. a, Representative image of a swarming motility assay with strains expressing wild-type or ΔN17 RhlR. Swarming assays were performed in technical duplicate with at least three biological replicates. **b,** Rhamnolipid production from the supernatants of strains expressing wild-type or ΔN17 RhlR as measured by Victoria Pure Blue BO and measuring absorbance of the dye at OD625 nm. Statistical significance was determined by unpaired t-test. **c,** C4-HSL was quantified in culture supernatants after 6 and 24 hours of growth by LC-MS using standard curve fitting. Data points represent 3 technical replicates from each of 2 biological replicates (6 points total per strain). Bars represent the mean; error bars represent SD. n.d., not detected (n.d.) denotes below the assay limit of detection. Given the limited number of biological replicates (n = 2), formal statistical comparisons were not performed. **d,** Same as in **c**, except for the quantification of pyocyanin.

**Figure S7.**
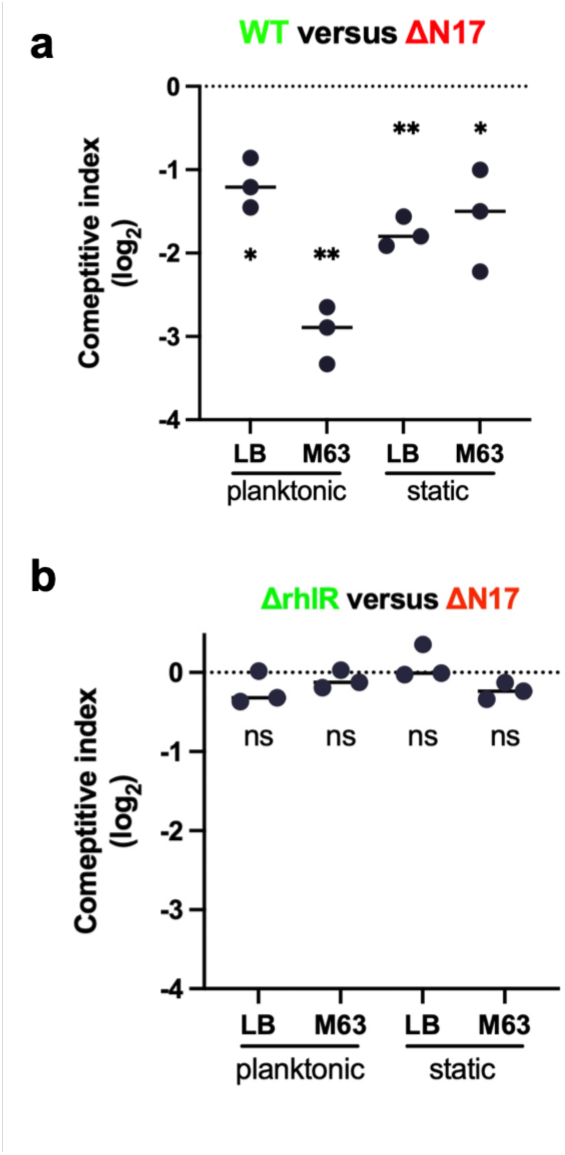
Stabilized RhlR imposes a competitive fitness cost that requires functional RhlR signaling. The reciprocal competitive fitness assays to Fig. 6a, comparing wild-type and ΔN17 RhlR strains across various growth conditions. Wild-type cells constitutively expressing mNeonGreen were mixed 1:1 with ΔN17 RhlR cells expressing mScarlet, grown for 24 hours under planktonic (LB and M63 media) or static (LB and M63 media) conditions, and enumerated by CFU plating. Data are represented as log₂ competitive index (CI), where negative values indicate fitness defects in ΔN17 relative to WT. Each point represents an independent biological replicate. Statistical significance was determined by one-sample t-test comparing log₂-CI values to 0 (no fitness defect); \*\**P* < 0.01; \*\*\**P* < 0.001.

**Figure S8.**
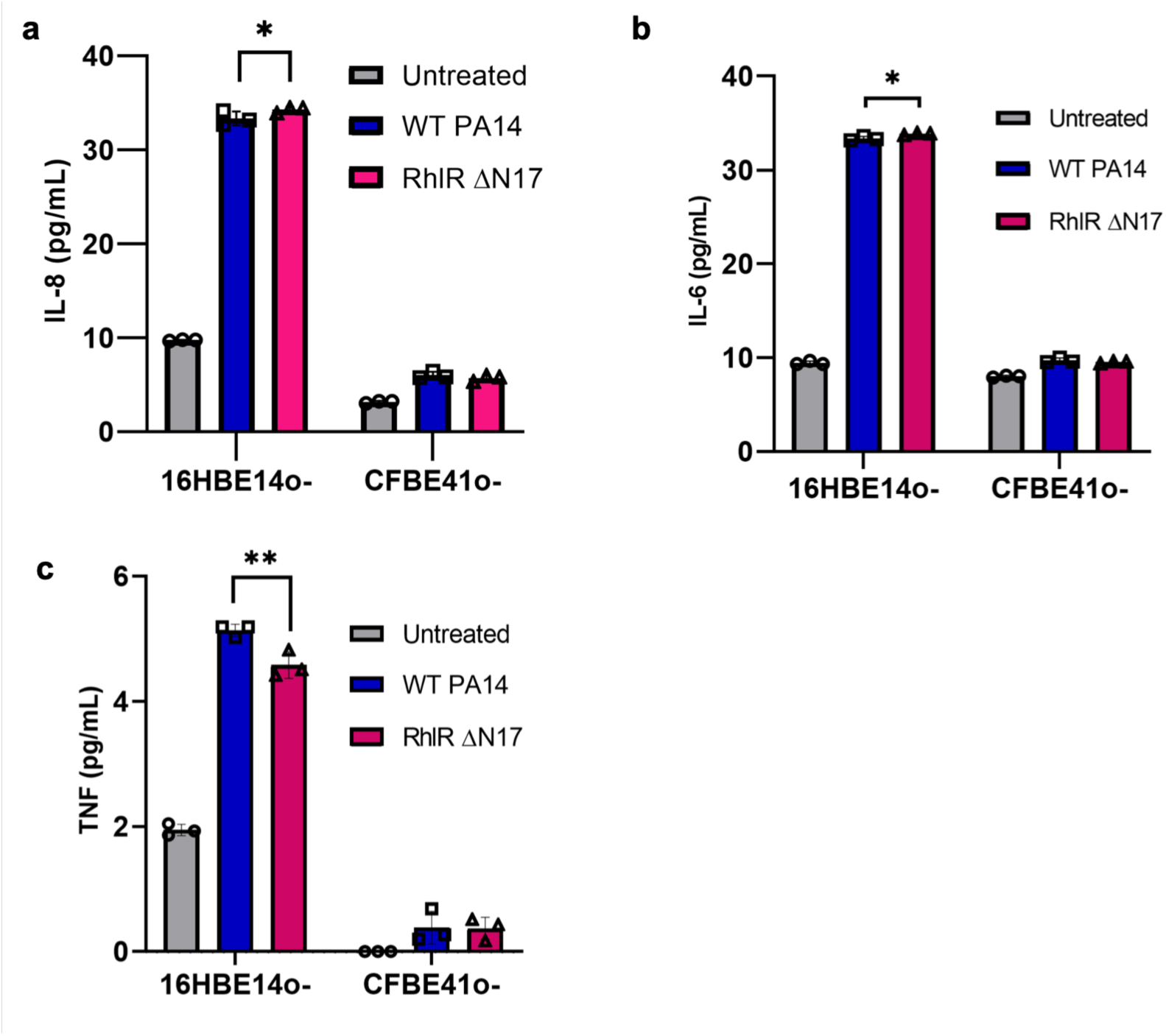
Cytokine production in lung epithelial cells is increased upon infection. IL-8 **(a)**, IL-6 **(b)**, and TNF **(c)** were measured in wild-type 16HBE14o- and CFTR F508del CFBE41o- bronchial epithelial cells in response to wild-type and ΔN17 RhlR via flow cytometry. Infections of the epithelial cells were done in biological triplicate. Statistical significance was determined by two-way ANOVA with Tukey’s multiple comparisons test; **(a)** \**P* = 0.0347, **(b)** \**P* = 0.0173, **(c)** \*\**P* = 0.0039.

## ACKNOWLEDGEMENTS

The authors thank all members of the Paczkowski lab as well as the research laboratories in the Division of Genetics at the Wadsworth Center for helpful discussions on the research, especially the Owen and Bartlett labs for shared equipment resources. We thank Dr. Joseph Wade (Wadsworth Center) for his advice on RNA-seq data processing. We thank Drs. Jia Jia Zhang and Tanya Baker (MIT) for the generous donation of strains. This work was made possible with the help of the dedicated staff scientists at the Advanced Genomics Technologies Cluster and the Media & Tissue Core facilities at the Wadsworth Center.

## FUNDING AND ADDITIONAL INFORMATION

This work was supported by National Institutes of Health training grant T32GM132066 and F31AI197475 to C.P.M., NIH grant R01GM14436101, New York Community Trust Foundation grant P19-000454, Cystic Fibrosis Foundation grant PACZKO21G0, and American Lung Association Innovation Award INALA2023 to J.E.P. The development of the cell culture model was supported by a SUNY Albany Grenander Award for Non-animal Methodologies in Research, Testing, and Education to M.L.S. and J.E.P. Cystic Fibrosis Foundation grants FERNAN24F0 supported S.E.F. and NAREN24R20 supported P.A.J.

